# Somatic mutation probabilities learned from the DNA sequence

**DOI:** 10.64898/2026.11.13.766380

**Authors:** Mario Aguilar-Herrador, Yana D. Müller, Melissa Sanabria, Anna R. Poetsch

**Author notes:** Equal contribution (Mario Aguilar-Herrador, Yana D. Müller).

## Abstract

Somatic mutations distribute heterogeneously over genomes, dependent on immediate sequence context, function, and non-genetic factors. Mutations present the foundation for somatic genome evolution towards age-related diseases.

Here, we present EAGLE-MUT, a group of DNA language models trained on sparse single base substitution data from esophageal adenocarcinoma (EAC) and melanoma (MELA) samples. In both cancer types, predicted mutation probabilities vary across several orders of magnitude, depending on sequence context. For example, T>G hotspots in EAC localize to A/T-rich ∼200 base pair DNA motifs, associated with nucleosome positioning. Coldspots localize to G/C-rich DNA, nucleosome linker DNA, CTCF binding sites, potential origins of replication, and intron-exon boundaries with a transcription strand bias. Fine-tuning EAGLE-MUT for individual patient samples leads to varying mutation distributions relative to genome elements and potential cancer driver mutations.

EAGLE-MUT provides new links between DNA sequence, genome function, and stability in individual samples, a foundation for understanding somatic genome evolution, and more personalized strategies for risk-assessment, prevention, and treatment.

## Introduction

Somatic mutations occur at distinct locations over the genome, shaped by their etiology, the physicochemical properties of the DNA sequence itself, the underlying epigenome^1,2^, gene expression^3^, replication timing^4^, and selective forces of somatic genome evolution. DNA strand specificity of mutagenesis is dependent on phasing^5^, strand-specific sequence content, direction of transcription and replication^6^. Genome regions prone to specific epigenetic regulation and strand-specific sequence patterns can be recognized by DNA language models^7^, which provides a link to the underlying DNA sequence and highlights strong connections between the DNA sequence and processes relevant for mutagenesis, such as DNA damage distribution^8^ and DNA replication timing^9^. The specific immediate sequence context is also a major determinant for occurrence of single base substitutions (SBS). Pattern differentiation of mutated 3-mers allows assignment of mutational signatures, which in many cases can be linked to an etiology of the underlying mutagenic mechanism^10–12^. Some mutagenic mechanisms are characterized by larger DNA sequence motifs, which can be linked to CCCTC-binding factor (CTCF) or transcription factor binding^13–15^, sequences prone to non-B-DNA structures^16–18^, and nucleosome coverage periodicity^19^.

Taken together, genome-wide distribution of mutations follows distinct patterns in somatic and cancer genomes of individuals. It builds the foundation for somatic evolutionary selection, thereby shaping potential somatic evolutionary trajectories. The extent to which the processes of somatic mutagenesis depend on larger DNA sequence patterns has remained elusive and presents a complex problem. Mutagenesis follows unique mechanisms, with every tissue and patient showing their own combination of interactive patterns. The outcome of mutagenesis, i.e. the mutations that can be measured, is sparse. Current models^20–25^ overcome sparsity by summarizing over genome elements or pooling mutations from different individual patient samples. This approach has been useful for defining sites of high mutation probabilities, i.e. mutation hotspots, over multiple patients^26^, yet it generalizes patterns of mutagenesis across individual tumour samples. Moreover, it does not allow us to find sites that are unlikely to mutate, i.e. coldspots.

Here we report a new group of models that generate per-nucleotide probabilities of somatic single base substitutions across the genome, based on a 500 base pair (bp) sequence context. The obtained mutation probabilities from esophageal adenocarcinoma (EAC) and melanoma (MELA) patient samples recapitulate known relationships with genome biology. Most strikingly, mutation probabilities for individual mutation types distribute relative to sequence motifs for nucleosome positions and reveal mutation coldspots at sites relevant for different genome biological function. Beyond pre-training with pooled mutations across a cohort, we also fine-tuned the general mutation patterns for individual samples, which shows strong local inter-individual differences with relevance for somatic genome evolution.

## Results

### EAGLE-MUT predicts per-nucleotide genome-wide mutation probabilities

To understand the impact of DNA sequence on somatic mutagenesis, we calculated a retrospective probability for every nucleotide in the genome to mutate *en route* to the observed mutation patterns. We trained a bidirectional recurrent neural network with long short-term memory (Bi-LSTM) in 500bp windows of the human reference genome (hg19) to convert the DNA sequence to its somatically evolved version with integrated single base substitutions (**Fig1A**). The model EAGLE-MUT (<u>E</u>fficient <u>A</u>nalysis with a <u>G</u>enome-wide <u>L</u>STM for <u>E</u>valuating <u>MUT</u>ation probabilities, “EAGLE” in the following) takes as input the one-hot-encoded DNA sequence of the reference genome and outputs probabilities for each nucleotide after somatic genome evolution with SBS. It thus converts the sparse distribution of SBS in somatic genomes into genome-wide per-nucleotide probabilities for SBS into each of the remaining three nucleotides. The information available to the model for training is therefore the sequence context of up to 500bp and the observed mutations, which reflect both mutagenic mechanisms by their immediate sequence context^10,12^, interactions of the mutagenic mechanisms with each other, and their interactions with the epigenome, where the epigenome is rooted in the DNA sequence, e.g. for DNA methylation-dependent mutations on CG dinucleotides^27,28^.

**Figure 1.**
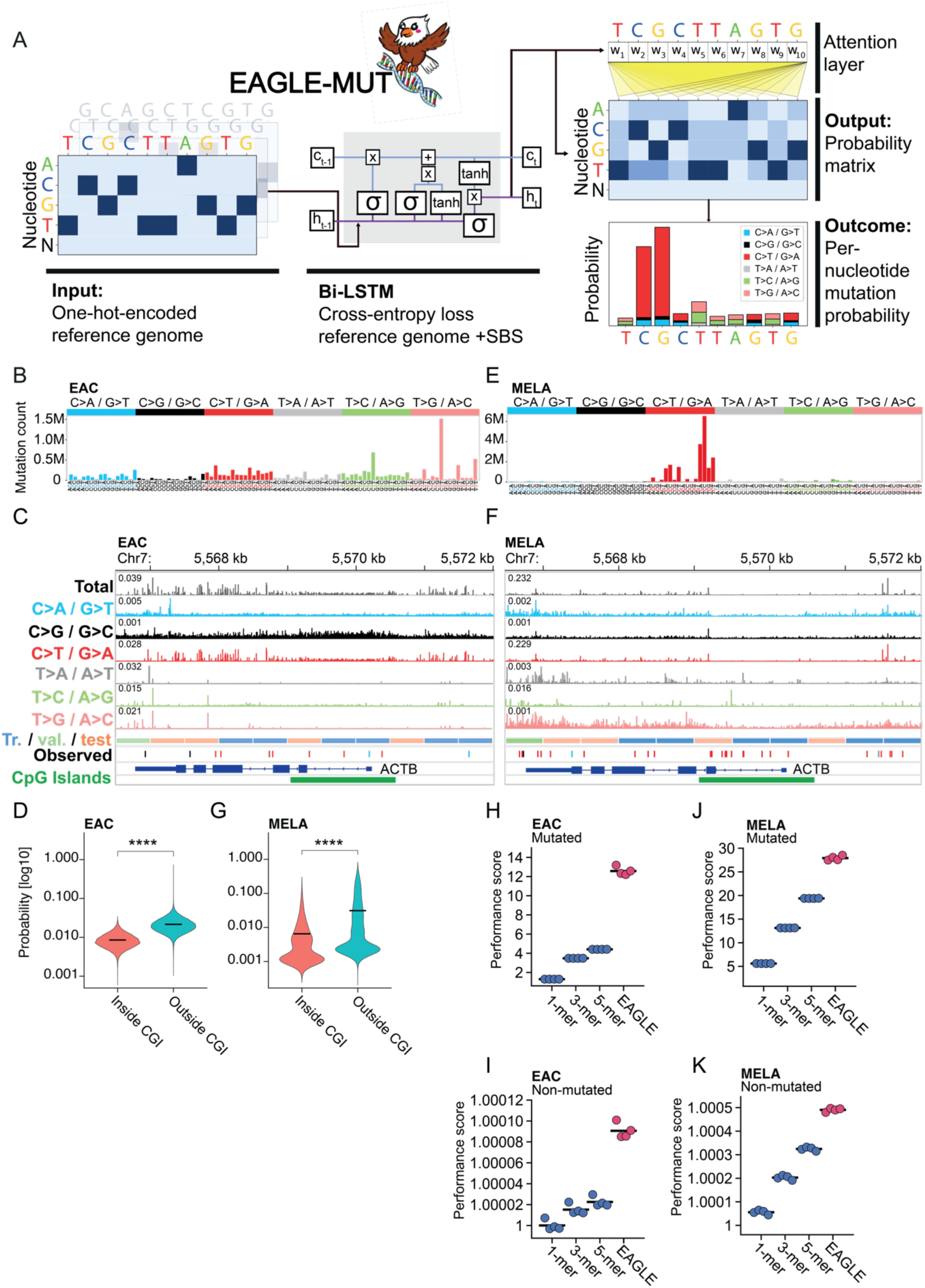
EAGLE-MUT learns genome-wide probabilities of single base substitutions. (A) Model design of EAGLE-MUT. EAGLE-MUT = <u>E</u>fficient <u>A</u>nalysis with <u>G</u>enome-wide <u>L</u>STM for <u>E</u>valuating <u>MUT</u>ation probabilities, Bi-LSTM = <u>Bi</u>directional recurrent neural network with <u>L</u>ong <u>S</u>hort-<u>T</u>erm <u>M</u>emory, SBS = <u>S</u>ingle <u>B</u>ase <u>S</u>ubstitution. (B) Total mutation counts over the cohort of esophageal adenocarcinoma, 423 samples from 409 patients, differentiated by mutation type in its trinucleotide context. (C) Relative mutation probability per mutation type over the *ACTB* locus. Probabilities range from 0 to the maximum value indicated on each track. (D) Probability assessment inside and outside of CpG islands for predicted CG>TG mutations in log10 scale for EAC. Bars indicate the mean, **** =p-value <0.0001; Wilcoxon test. (E) Total mutation counts over the cohort of skin melanoma, 199 samples from 183 patients, differentiated by mutation types in their trinucleotide context. (F) Relative mutation probability per mutation type over the *ACTB* locus for MELA. Probabilities range from 0 to the maximum value indicated on each track. (G) Probability assessment inside and outside of CpG islands for predicted CG>TG mutations in log10 scale for MELA. Bars indicate the mean, **** =p-value <0.0001; Wilcoxon test. (H) Performance assessment as the ratio of EAGLE-predicted probability in the test set at sites of observed mutations (Mutated) in EAC versus a random model, in comparison to proportional assignment of probabilities based on the observed mutations in the associated k-mer context. 4 models are shown, trained with 4 different splits of training, validation, and test sets. Bars indicate the mean. (I) Performance assessment of sites without observed mutations (Non-mutated) in EAC, analogous to (H). (J) Performance assessment as the ratio of EAGLE-predicted probability in the test set at sites of observed mutations (Mutated) in MELA versus a random model, in comparison to proportional assignment of probabilities based on the observed mutations in the associated k-mer context. 4 models are shown, trained with 4 different splits of training, validation, and test sets. Bars indicate the mean. (K) Performance assessment of sites without observed mutations (Non-mutated) in EAC, analogous to (J). EAC = Esophageal Adenocarcinoma, MELA = Melanoma, Tr. = Training set, val. = Validation set, test = Test set.

EAGLE was initially trained on SBS pooled from 423 genomes (409 patients) from the ESAD EAC cohort of the PCAWG project^29^. The 12,376,220 SBS from the pooled mutation catalogues are characterised by high contributions of T>G/A>C transversions, of which CTT>CGT (n=1,518,274) is the most frequent (**Fig1B**). By calling mutation signatures on the cohort with SigProfilerExtractor^30^, ten mutation signatures were derived (**FigS1A**). Most of them are related to known mutation signatures described for this cohort in PCAWG^11^, except for SBS100, which has been described independently^31^. Of note, two of the signatures are related to COSMIC SBS17, and one displayed a higher relative contribution of T>G transversions. These signatures can be linked to gastro-esophageal reflux^6^. We clustered the signatures using k-means clustering and assigned five patient sample groups (**FigS1B**). Groups i-iii are characterized by a combination of “background” signatures and gastro-esophageal reflux. They are assigned to three different clusters according to their signature balance. Group iv shows mutation profiles linked to APOBEC activity, and group v shows defects in replicative proofreading and mismatch repair.

Across a typical house-keeping gene, *ACTB*, a region that contains 11 observed SBS, the distribution of EAGLE-imputed mutation probabilities is heterogeneous and differs by mutation type (**Fig1C**). Taking all six mutation types together, EAGLE’s mutation probability predictions show large local differences with a spread of probability over several orders of magnitude. C>A and C>G show largely smooth distributions of probabilities with some local peaks. This is in contrast to C>T, which occurs predominantly in a CG context. However, within CpG islands, where CGs are denser and thus more locations show elevated mutation probability, probabilities for each CG are reduced, an observation that scales genome-wide with CG dinucleotides (**Fig1D**). While the mean probability of CG>TG in CpG islands is 0.0085, it is 4-fold increased to 0.021 for CG dinucleotides outside CpG islands (p<0.0001 Wilcoxon test). The mutagenic process on CG dinucleotides is dependent on DNA methylation^27,28^, which is generally lower in promoter CpG islands in healthy tissue^32^. EAGLE recognizes this relationship with the epigenome and assigns the probabilities accordingly. The 3’-UTR of *ACTB* also shows a spike of T>A. This mutation occurs on an A_n_T_n_ stretch. The training data contain multiple variants in this sequence context, which also have the potential to be the result of sequencing or variant calling errors. Lastly, T>C and T>G show very heterogeneous probabilities with clear cold- and hotspots.

Complementarily, we trained EAGLE on SBS (n=24,579,703) from 199 MELA samples of 183 patients. This tumour type is dominated by C>T mutations from pyrimidine dimers with the most frequent SBSs occurring in a TCC context (n=6,752,406; **Fig1E**), which predominantly shape the predictions in the *ACTB* locus (**Fig1F**). Also in MELA, CG>TG is predicted as less likely in CpG islands than outside (mean probability = 0,006 vs. 0,031, p<0.0001, Wilcoxon test; **Fig1G**).

### EAGLE recapitulates observed mutation patterns

To benchmark the performance of EAGLE, we used k-mer context models, which represent the immediate sequence context and typically serve to estimate selection-independent mutation backgrounds (**Fig1H-K, FigS2&3**). We treated the probability of mutations and non-mutations, i.e. sites without an observed mutation, as two separate tasks, which we evaluated in the genomic regions that encompass the test set. The performance score represents the average ratio of the model’s prediction over a null model that assumes a uniform and random distribution of mutations across the genome and therefore allows us to assess the relative change of performance with increasing sequence context. Increase of sequence context leads to a steady improvement of performance. EAGLE predicts mutations and non-mutation in EAC 12.6- and 1.00009-fold over random, respectively. In MELA, EAGLE predicts mutations and non-mutation 27.9- and 1.00049-fold over random, respectively. Interestingly, the gradual increase of sequence context indicates less relevance for the immediate sequence context for EAC given that EAGLE highly outperforms the k-mer models. MELA samples, however, show a substantial performance increase with including the information of the mutated base (1-mer) and increasingly so with the sequence context of increasingly large k-mers. This indicates that for the mutagenic mechanisms in EAC a larger sequence context is highly relevant, whereas for MELA the immediate sequence context is a major factor for mutation distribution and the relative benefit of the additional EAGLE context is smaller. Similar relative performances can be derived with alternative methods that interrogate Pseudo R^2^ and variance explained (**FigS2**). Increasing the k-mer context beyond 5-mers marginally improves performance, but is increasingly affected by sparsity of some k-mers in the genome (**FigS3A-G**). Varying the sequence context of EAGLE (**FigS3H&I**) results in inferior performance below 500bp and no steady increase beyond, which lets us conclude that with this architecture, 500bp windows lead to the smallest model with optimal performance.

Next we interrogated EAGLE performance relative to the SBSs’ trinucleotide context. Mean mutation probability predictions by trinucleotide show a cosine similarity with the observed mutation rate of 0.998 and 0.999 for EAC and MELA, respectively. In EAC, EAGLE predictions show a Spearman’s R of 0.94 (p<0.001) as compared to the 3-mer models (**Fig2A**) and Spearman’s R of 0.96 (p<0.001) in MELA (**Fig2B**). This shows how EAGLE recapitulates mutation patterns by differentiating the underlying 3-mer context. Interestingly, EAGLE predictions differ substantially within the 3-mer context (**Fig2C&E**), spanning up to 5.2 orders of magnitude for the most frequent mutation type in EAC, CTT>CGT, and 3.9 orders of magnitude for TCC>TTC in MELA. This suggests that EAGLE is able to learn context well beyond 3-mers. However, not every mutation type is learned equally well (**Fig2D&F**) due to their different frequency and underlying biology. Differentiating between mutation types, the mutation prediction performance peaks at a score of 114.2 for TTA>TAA and 56.6 for CTT>CGT in EAC and in MELA at a score of 80.0 for TCG>TTG and 62.0 for TCC>TTC.

**Figure 2.**
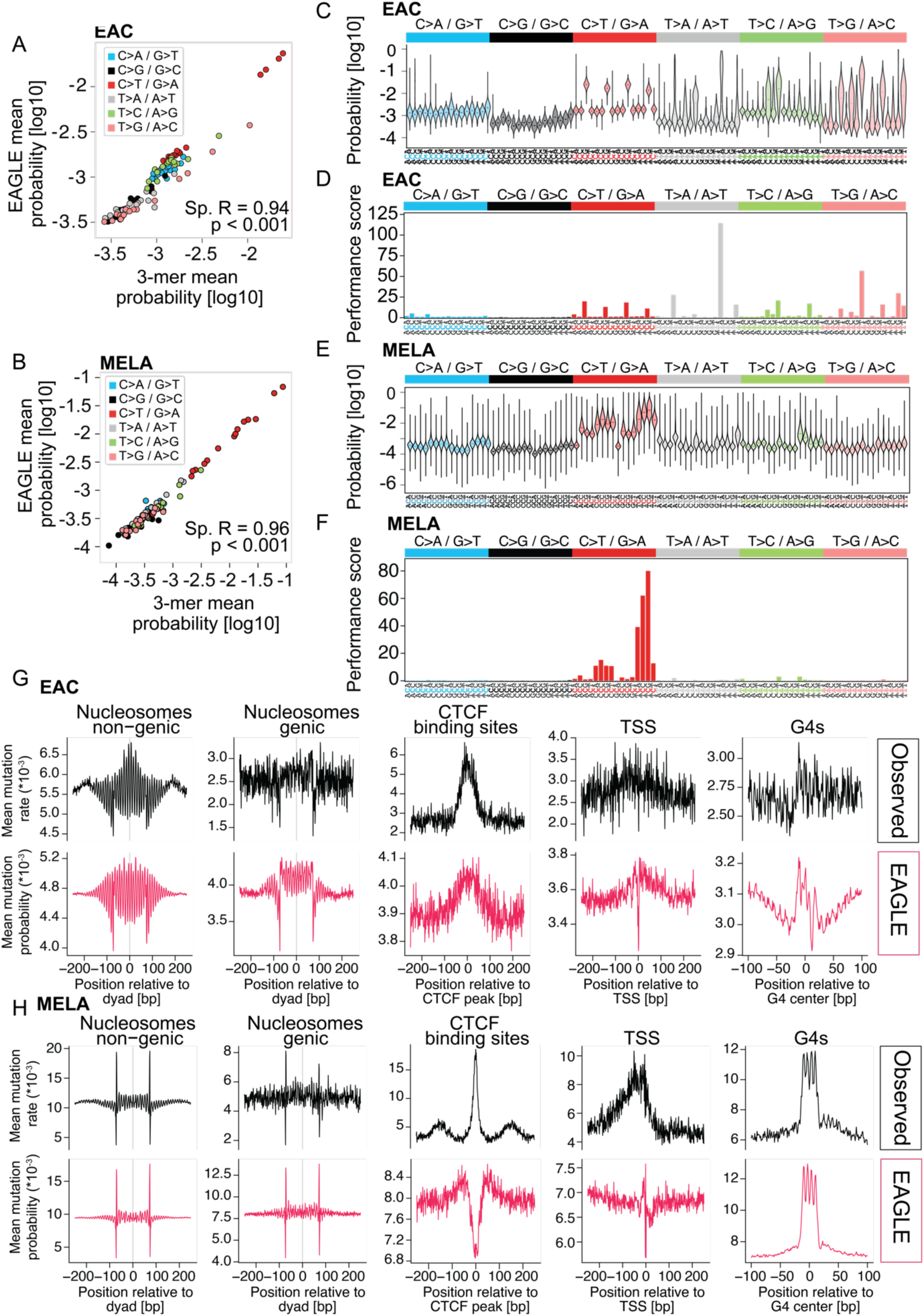
EAGLE recapitulates known mutation patterns. (A) Correlation of mean probabilities based on mutation distribution in their 3-mer context, compared to mean EAC EAGLE prediction. Sp.R = Spearman’s R. The p-value was derived through a Spearman’s test with an asymptotic t approximation. (B) Correlation of mean probabilities based on mutation distribution in their 3-mer context, compared to mean MELA EAGLE prediction. Sp.R = Spearman’s R. The p-value was derived through a Spearman’s test with an asymptotic t approximation. . (C) Distribution of EAC EAGLE predictions for each mutation type in its trinucleotide context. The horizontal line indicates the mean. (D) Performance score of EAC EAGLE mutation prediction for each mutation type in its trinucleotide context. (E) Distribution of MELA EAGLE predictions for each mutation type in its trinucleotide context. The horizontal line indicates the mean. (F) Performance score of MELA EAGLE mutation prediction for each mutation type in its trinucleotide context. (G) and (H) Distribution of mean EAGLE predictions of mutation probability in comparison to mean observed SBS rate distribution, relative to genome elements. TSS = Transcriptional Start Site, CTCF = CCCTC-Binding Factor, G4s = predicted G-quadruplex structures.

To interrogate the larger sequence context learned, we compared mean EAGLE predictions against mean observed mutation rates and k-mer context models over genome elements (**Fig2G&H**, **FigS4&5**). Around non-genic nucleosomes, SBS in both EAC and MELA accumulate relative to the orientation of the minor groove to the nucleosome^19^, yet with different phasing and distinct patterns around the linker DNA. EAGLE predictions recapitulate the periodicity of the minor groove. For genic nucleosomes, observed mutations are sparse, leading to noisy profiles. EAGLE overcomes this sparsity and presents a periodic pattern that also indicates the orientation of the minor groove. Absolute values differ between the predicted and observed, because the range of the model leads to a focus on local patterns rather than long-range relationships with chromatin architecture, which affects the base line of the predictions. Interestingly, while in EAC EAGLE recapitulates the peak of the observed mutation profile over CTCF binding sites, yet with a smaller range as compared to the flanking region, it fails to recapitulate mutation patterns at CTCF binding sites for MELA. Intrigued by this mismatch, we investigated further, why the models struggle with this specific task (**Fig. S6**) and found a similar mismatch to ETS1 binding sites, which are also expected mutation hotspots^33^. For both protein binding sites we find that EAGLE underestimates the mutation rates in the bound motifs and overestimates mutation rates in the higher number of unbound motifs (**FigS6A**). However, when differentiating the motifs by whether they are observed to be mutated, a locally increased prediction of mutation probability can be observed (**FigS6B**) and sites with increased probabilities are more frequently mutated (NES_ETS1_=1.52, p_ETS1_=2.7e-14, NES_CTCF_=1.61, p_CTCF_=1.2e-93; **FigS6C**). Therefore, the underestimate of EAGLE of these sites is likely due to the imbalance of mutated versus unmutated motifs, where EAGLE regresses to the mean of the mutation probability. Potential consequences of positive selection, which would not be recognizable to EAGLE, may add to the disagreement between observed and predicted mutation densities. Transcriptional start and termination sites (TSS and TTS), exon junctions, and Alu retrotransposons (**Fig2G&H**, **FigS4&5**) also show distinct mutation probability patterns that approximate the observed mutation patterns with underestimates at sites of expected CTCF and transcription factor binding in MELA. In EAC, mutation profiles on predicted G-quadruplex structures (G4s) are difficult to model accurately with k-mers, nonetheless EAGLE indicates a distinct asymmetric mutation pattern.

We therefore conclude that EAGLE learns profiles beyond the 3-mer pattern of mutagenesis and uses a larger sequence context to largely recapitulate mutation profiles of the ground truth appropriately along genome elements with distinct patterns of mutagenesis.

### Nucleosome positioning motifs are the main determinants for CTT>CGT mutagenesis in EAC

The imputation of mutation probability overcomes sparsity of observed mutations and thus allows us to investigate specific mutation types in more detail. First, we focus on CTT>CGT, the most frequent context for SBS in EAC, a mutation type related to gastro-esophageal reflux. Linked to the COSMIC-mutation signature SBS17^10^, T>G mutations are believed to be the result of mismatches by erroneous incorporation of 8-oxo-guanidine from the nucleotide pool opposite adenine^34^. This DNA lesion - mutation pair is known to be distributed heterogeneously over the genome^35,36^. When investigating CTT>CGT in the context of genome elements (**Fig3A**, **FigS7A**), patterns observed for this mutation type in the non-genic nucleosomes, G4s, Alu retrotransposons, and exon boundaries are largely recapitulated by EAGLE. Genic nucleosomes, TSS, TTS, G4s, and exon boundaries suffer from sparsity in the ground truth, yet EAGLE indicates a more differentiated pattern of mutation probabilities, including periodic mutation probabilities relative to genic nucleosomes, which are hardly discernible in the profiles of observed mutations. Interestingly, for the profile for CTCF binding sites, EAGLE predicts a dip, where the ground truth indicates a peak, likely for analogous reasons to the predicted MELA mutation rate distributions over these sites (see above).

**Figure 3.**
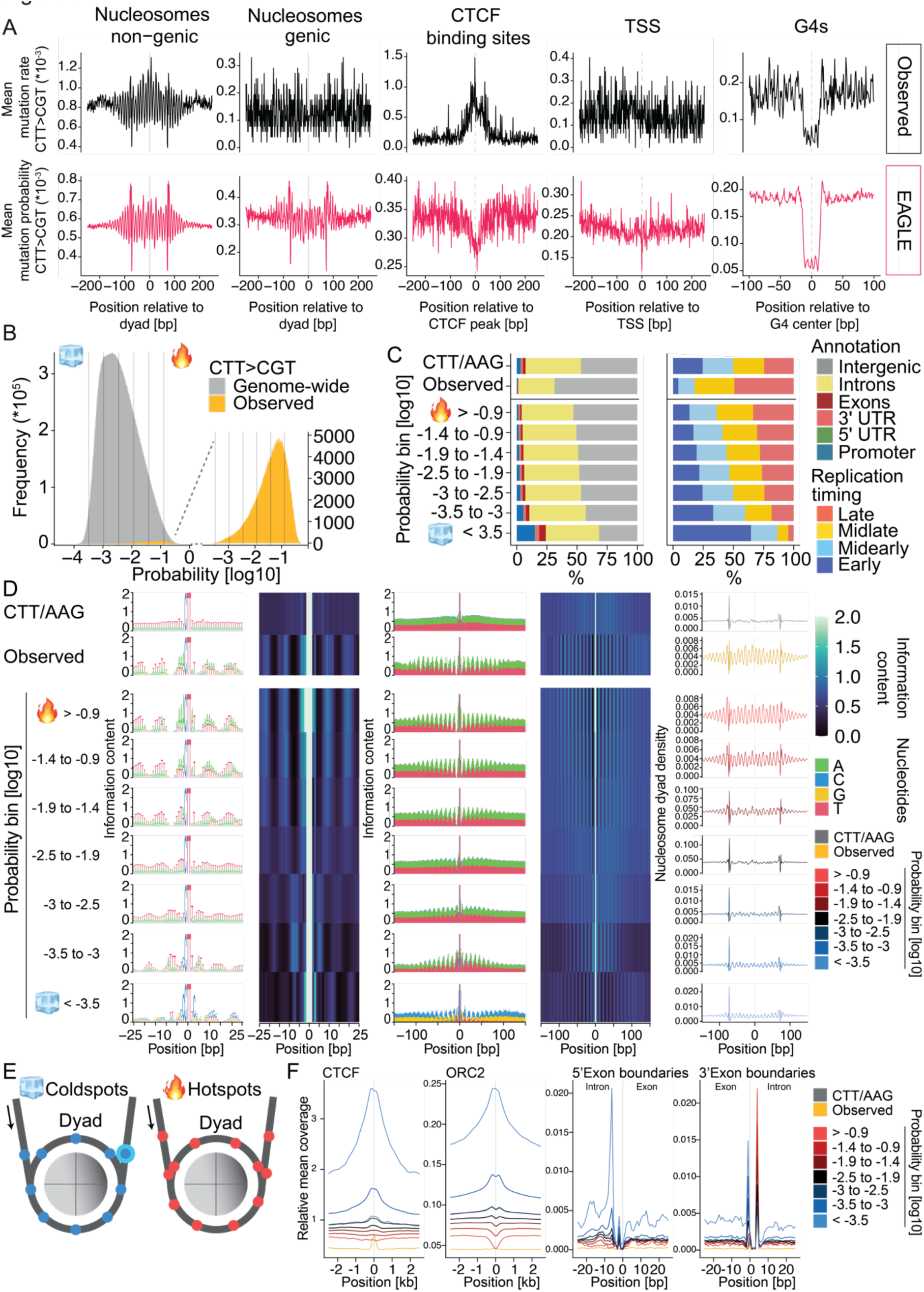
CTT>CGT mutation probabilities in EAC follow distinct sequence patterns related to genome biology. (A) Mean mutation rate distribution of EAGLE predictions for CTT>CGT/AAG>ACG in comparison to mean observed SBS distribution, relative to genome elements. TSS = Transcriptional Start Site, CTCF = CCCTC-Binding Factor, G4s = predicted G-quadruplex structures. (B) Distribution of EAGLE predictions on CTT>CGT/AAG>ACG genome wide and at sites of observed mutations. Ice cube and flame symbols denote the bins with the extreme (1%) low and high probability, classified as cold- and hotspots, respectively. (C) Gene- and DNA replication timing-dependent annotation of CTT>CGT/AAG>ACG with different mutation probabilities in comparison to observed mutations and background distribution of CTT/AAG. (D) Sequence motif patterns on CTT>CGT/AAG>ACG with different mutation probabilities in comparison to observed mutations and background distribution of CTT/AAG. Information content represents the differential of 2 bits and the information entropy of the sequences flanking the mutated trinucleotide, resolved by nucleotide as a motif visualization and as an information content summary. Nucleosome dyad density represents the mean proportion of nucleosome dyads irrespective of their relationship with genes, relative to the trinucleotide. (E) Schematic for the location of mutation cold- and hotspots relative to nucleosome dyads. (F) Relative mean coverage of CTCF, ORC2, and exon boundaries on CTT>CGT/AAG>ACG with different mutation probabilities in comparison to observed mutations and background distribution of CTT/AAG.

CTT>CGT mutation rates range over 5.2 orders of magnitude with a longer tail towards higher mutation probabilities (**Fig3B**). The peak of predicted log10 probabilities is at -2.72, the peak of log10 prediction at the sites of observed mutations is at -1.08, with an equal spread, yet leaning towards higher probabilities and a longer tail towards lower probabilities. Next, we grouped the genome-wide probabilities of CTT>CGT mutations into seven categories to investigate relationships with sequence and genome biology. First, we interrogated the mutation probability of these trinucleotides with respect to gene annotation (**Fig3C**). The majority of the underlying trinucleotides lie in intronic and intergenic regions (45.9% and 46.8% respectively), yet with a balanced distribution relative to replication timing. The observed mutations are enriched in intergenic regions (68.7%), and in mid-late and late replicating DNA (32.9% and 49.2%, respectively). CTT/AAG sites with higher mutation probability follow the trend, whereas sites with lower mutation probabilities are increasingly detected in coding sequence, untranslated regions (UTRs), promoters, as well as early replicating DNA. This relationship to gene annotation and replication timing has already been described by Tomkova *et al.*^6^, yet the predictions solely based on DNA sequence data indicate that this relationship is partially rooted in the sequence. Similarly, the known transcriptional strand bias of CTT>CGT mutagenesis can be recapitulated (**FigS7B&C**). However, replication strand bias display an odds ratio of 1.066 ± 0.005 of the observed mutated trinucleotides relative to trinucleotide distribution andthe opposite is observed for the trinucleotides of most likely probability for CTT>CGT (0.944 ± 0.005). This is likely due to sequence-independent mechanisms or the 500bp training window size of EAGLE focusing on local sequence context, insufficiently broad for the requirements for replication strand and timing.

Mutation profiles follow distinct long-range motifs (**Fig3D**), reflected by the sites of predicted high probability, i.e. hotspots. They are composed of an asymmetric short-range motif (WWW<u>CTT</u>MW) and A/T rich 7-mer waves spaced by 3bp. The range of these motifs exceeds 200bp, which shows that EAGLE can indeed learn both short- and intermediate-range relationships between sequence and mutation probability. With decreasing probability, the repetitive motif increasingly fades out, being re-established at probabilities <10^-3^ with alternate phasing when compared to the higher probabilities. Coldspots also follow a distinct asymmetric motif (BC<u>CTT</u>BC), which is preceded by a TT motif and flanked by a GC-rich wave pattern analogous to the hotspots. These motifs relate to the orientation of the DNA’s minor groove relative to the positioning of nucleosomes. Pich *et al.* have reported that these mutations localize preferentially at the minor groove facing away from the nucleosome^19^. Inversely, we show that relative to the mutations, nucleosome dyads are positioned periodically relative to the mutation hotspots (**Fig3D&E**). Complementarily, nucleosome dyads are enriched on coldspots and in periodical positions indicating sequence content that facilitates the minor groove pointing towards the nucleosomes. Interestingly, there is an enrichment of dyads about 73bp in the 5’ direction of the coldspots, which is already present in the background distribution of CTT/AAG. This position is indicative of the linker DNA 3’ of the nucleosome position and suggestive of a protective mechanism from mutagenesis, which could be due to increased activity of 8-oxoguanine DNA glycosylase-1 (OGG1)^37^. While the nucleosome positioning sequence was identified as a major factor for mutation probabilities, the linker DNA signal explains only a fraction of the coldspots. We thus interrogated other genome biological features from G/C rich DNA in their relationship with mutation probabilities, focusing on CTCF, ORC2 as a marker of potential origins of replication, and exon boundaries (**Fig3F**). Surprisingly, the general signal of all three is broadly elevated relative to the cold- and hotspots, yet with a local enrichment relative to the coldspots. This indicates that regions of low mutation rates can be attributed to several mechanisms, which prompted us to describe coldspots in more detail.

### EAGLE identifies mutation coldspots in EAC with different connections to genome function

We have identified five types of CTT>CGT coldspots linked to gene promoters, CTCF binding sites, potential origins of replication, linker DNA, and exon boundaries. These coldspots are largely mutually exclusive (**Fig4A**). We then investigated the relationship to transcription strand bias and confirmed largely the association with CTT mutations on the transcribed strand, except for coldspots associated with exon boundaries (**Fig4B**). The 5’ exon boundary is biased to the transcribed strand with an odds ratio of 2.17 ± 0.05, yet coldspots relative to the 3’ exon boundary are enriched in the non-transcribed strand with an odds ratio of 0.78 ± 0.02. Indeed, exon boundaries in sense or antisense to the coldspots show distinct profiles (**Fig4C**). That EAGLE learns a relationship with transcription points to a sequence-mediated mechanism, acting either directly or via transcription-coupled damage or repair. Given the distinct biological origins of coldspots, we differentiated the underlying motifs (**Fig4D**). While they all follow a similar periodic GC-rich wave pattern, they also show specific differences. Coldspots within the 3’ linker DNA, 73 ± 1bp or 76 ± 1bp from a nucleosome dyad, show a clear asymmetric motif with a local asymmetry between As and Ts. The position of the 5’ linker DNA shows a break of the G/C richness and a distinctive motif. These characteristics are different from coldspots associated with CTCF, ORC2, and promoters. Coldspots on exon boundaries show an asymmetric pattern with a CG dinucleotide enrichment within the location of exons. The intron side displays a G/C-rich wave pattern similar to the sequence foundation of nucleosome binding. There are a number of coldspots that could not be assigned to a group. A possible explanation might be a sequence associated with biological features that may be present while not being used in the tissue from which the data has been obtained. It is thus likely for this group to originate from a mixture of the described subgroups or, indeed, yet unidentified biological features. In conclusion, we can show that coldspots of mutagenesis lie at biologically relevant locations in the genome, a relationship between sequence and its stability to have evolved as a protection mechanism for such locations where mutation may have substantial genome-functional consequences.

**Figure 4.**
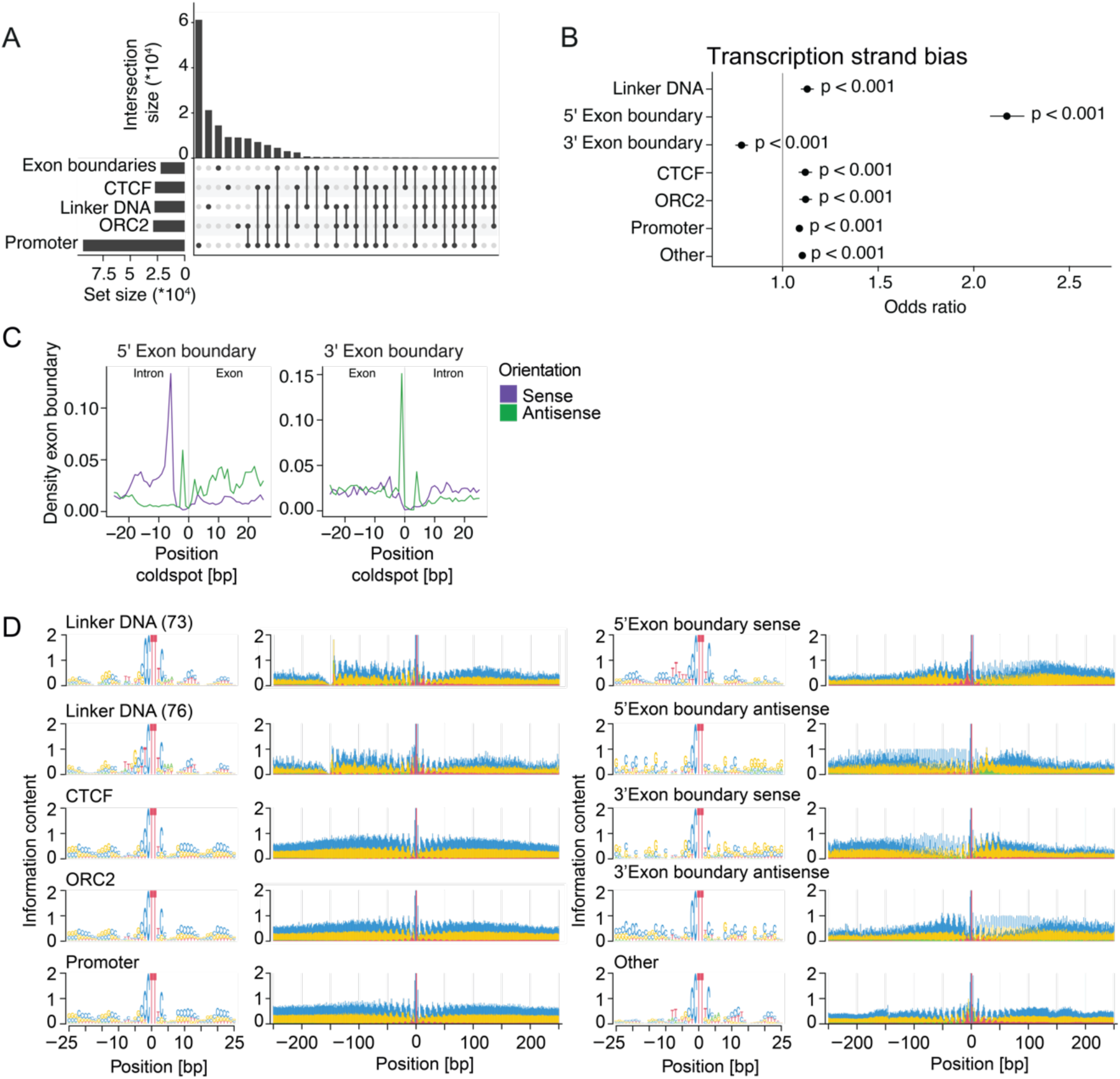
CTT>CGT coldspots in EAC localize at different sites of genome biological importance. (A) Comparison of annotating CTT>CGT/AAG>ACG sites defined as coldspots. (B) Transcription strand bias of EAGLE predictions for different CTT>CGT/AAG>ACG coldspots. Odds ratios are calculated with associated p-values based on the imbalance of the strands, corrected for the imbalance of the underlying trinucleotide. (C) Differentiation of CTT>CGT/AAG>ACG coldspots localized at exon boundaries by orientation and location. (D) Sequence motif patterns on different CTT>CGT/AAG>ACG coldspots. Information content represents the differential of 2 bits and the information entropy of the sequences flanking the mutated trinucleotide, resolved by nucleotide as a motif visualization.

CCT>CTT mutations in MELA are predicted with transcription strand bias and follow nucleosome positioning

Analogously we investigated, as the dominant mutation type, CCT>CTT mutations in MELA in more detail. The mutation distribution profiles over genome features largely reflect the total mutation rates (**FigS8A**, **Fig2H)**. The predicted mutation probability ranges 3.9 orders of magnitude (**Fig5A**) with an enrichment of hotspots in non-genic and late-replicating parts of the genome (**Fig5B**). Coldspots are enriched in promoters and early replicating DNA. Interestingly, also this mutation type follows distinct sequence contexts that relate to nucleosome positioning (**Fig5C**). Hotspots are characterized by TT<u>TCC</u>TTT with A/T-rich waves that relate to the helix’ position relative to nucleosomes and suggest hotspots on the nucleosome dyad. Predicted coldspots show the alternate phasing of nucleosome positioning in comparison to the hotspots. Enriched in GC-rich DNA with 38.4% in CpG islands and an asymmetric motif (Y<u>TCC</u>AY), they also show an asymmetric pattern at the linker DNA. The positioning relative to CTCF, ETS1 binding sites, potential origins of replication, and exon boundaries (**Fig5D**) shows an enrichment particularly at the coldspots, analogous to CTT>CGT mutations in EAC. Mutations from pyrimidine dimers in melanoma follow a strong transcription strand bias^38^. We therefore investigated whether we can observe this bias in the EAGLE predictions (**Fig5E, FigS8B**). Indeed, the predicted hotspots follow the observed transcription and replication strand biases, while the coldspots show the opposite relationship. This indicates that the sequence content indicates transcription and replication direction, so that this relationship is recognizable to the model by the sequence. Also foundation DNA language models like GROVER recognize this directionality without additional data but the DNA sequence^7^. Therefore, it is to be expected that this relationship is at least partially recognizable by the sequence, which we observe here as being stronger for TCC>TTC in MELA than the CTT>CGT in EAC.

**Figure 5.**
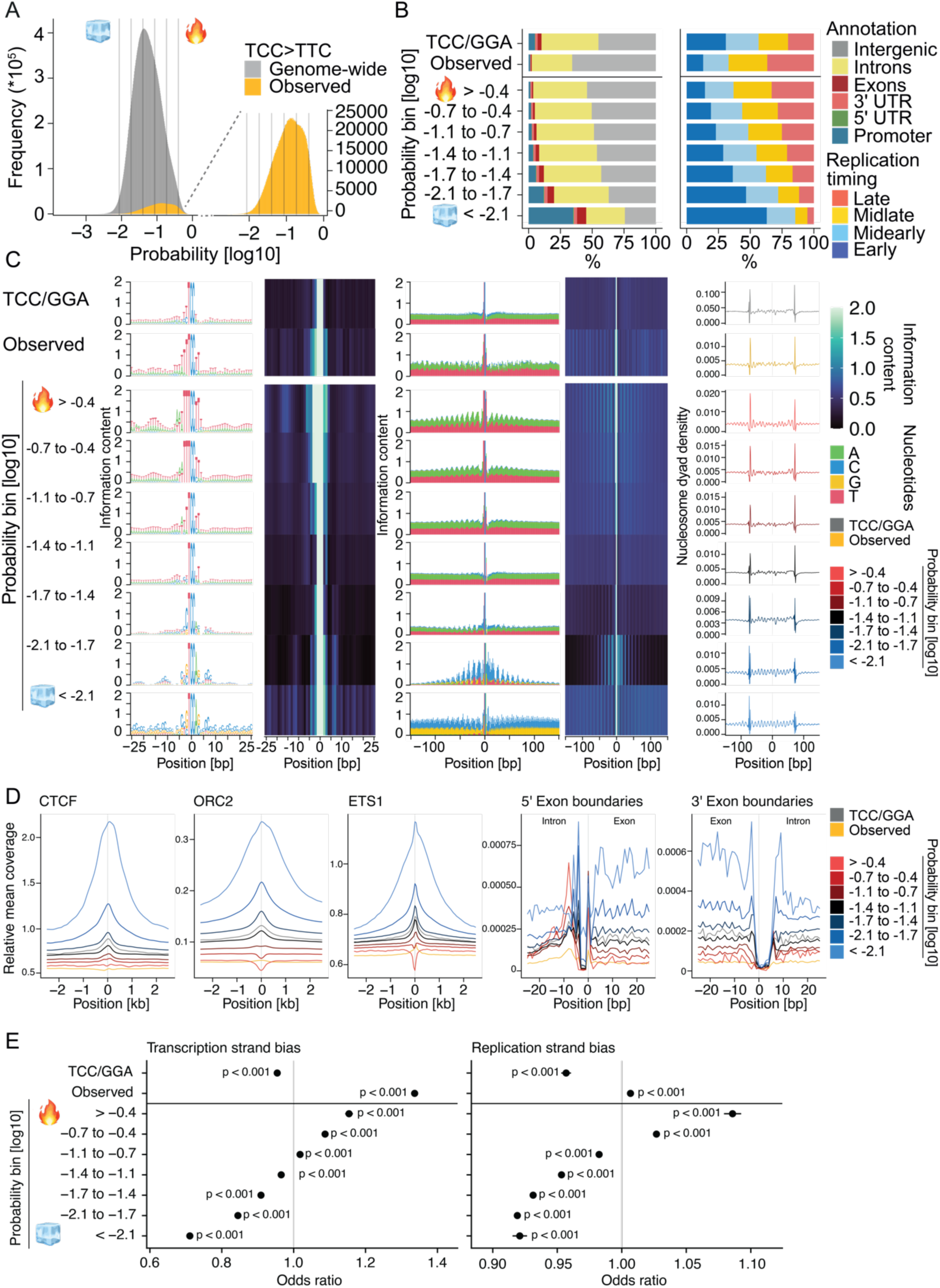
TCC>TTC mutation probabilities in MELA follow distinct sequence patterns related to genome biology. (A) Distribution of EAGLE predictions on TCC>TTC/GGA>GAA genome wide and at sites of observed mutations. Ice cube and flame symbols denote bins classified as cold- and hotspots, respectively. (B) Gene- and DNA replication timing-dependent annotation of TCC>TTC/GGA>GAA with different mutation probabilities in comparison to observed mutations and background distribution of TCC/GGA. (C) Sequence motif patterns on TCC>TTC/GGA>GAA with different mutation probabilities in comparison to observed mutations and background distribution of TCC/GGA. Information content represents the differential of 2 bits and the information entropy of the sequences flanking the mutated trinucleotide, resolved by nucleotide as a motif visualization and as an information content summary. Nucleosome dyad density represents the mean proportion of nucleosome dyads irrespective of their relationship with genes, relative to the trinucleotide. (D) Relative mean coverage of CTCF, ORC2, ETS1, and exon boundaries on TCC>TTC/GGA>GAA with different mutation probabilities in comparison to observed mutations and background distribution of TCC/GGA. (E) Transcription- and DNA replication strand bias of EAGLE predictions for TCC>TTC/GGA>GAA in comparison to mean observed SBS distribution and background distribution of TCC/GGA. Odds ratios are calculated with associated p-values based on the imbalance of the strands.

### Mutation cold- and hotspots in EAC and MELA follow similar patterns throughout mutation types

To generalize from the specific mutation sites, we explored all 96 mutation types of SBS in the trinucleotide context. We annotated cold- and hotspots relative to the underlying sequence information content (**FigS9A&10A, FigS11-14**), nucleosome binding (**FigS9B&10B**), genes and replication timing (**FigS9C&10C**), exon boundaries (**FigS9D&10D**), CTCF, binding (**FigS9E&10E**), potential origins of replication (**FigS9F&10F**), and for MELA, ETS1 binding (**FigS10G**). Generally, the coldspots of all mutation types tend to localize preferentially to early replicating DNA, promoters and genes, while hotspots prefer intronic and intergenic regions, more pronounced than the observed mutations. Interestingly, some coldspots for C>T and C>A with a following G show a tendency towards promoter sequence, which is not reflected in replication timing preference. Indeed, exon boundaries, CTCF, ORC2, ETS1 (for MELA), and linker DNA are enriched on the coldspots of several mutation types. For CTCF and ORC2 binding, C>A and C>T mutations followed by a G are the most prevalent ones. Both cold- and hotspots exhibit distinct motifs in the immediate sequence context that are more pronounced than for the observed mutations (**FigS11&13)**. Interestingly, the repetitive pattern for observed hotspots are apparent for many SBS types in MELA and EAC (**FigS12&14)**. Aligned with the enrichment in promoters and coding sequence, coldspots of most mutation types have transcriptional strand bias (**FigS15A&C**), in EAC from an odds ratio of 0.69 ± 0.01 for ACA>ATA to 1.33 ± 0.01 for GCG>GAG, in MELA from an odds ratio of 0.67 ± 0.01 for ATA>AGA to 1.82 ± 0.02 for GCG>GAG. Most mutation types display replication strand bias for predicted coldspots and hotspots (**FigS15B&D**). In conclusion, we are able to show that EAGLE learns relationships between mutation probabilities and genome biology.

### Genome-wide mutation probabilities are individually different

Previous analyses were performed on a pool of SBS from multiple patients, which raises the question of how consistent such patterns might be between individual cancer genomes. Therefore, we employed a pre-training/fine-tuning strategy with the model pre-trained on SBS pooled from the entire cohort for EAC and MELA and fine-tuned on individual samples. Assessing performance, we find that this strategy leads to a better outcome than either strategy alone (**Fig6A&B**). When applying the model trained on the entire cohort on individual samples, we reach a median performance score of 12.9 for predicting mutations correctly in EAC, and 26.7 in MELA. Training an EAGLE model for each sample individually, a lower median performance is reached (11.3 in EAC and 13.0 in MELA). Interestingly, the latter strategy is favorable for about 40% of the samples in EAC, yet inferior for the vast majority in MELA. Combining both training strategies through pre-training and fine-tuning, we generally achieve the highest performance with a median of 18.4 in EAC and 29.8 in MELA, which also exceeds the k-mer models (**FigS16A&C**). It would be expected that more mutations per sample would have led to improved performance. Even though the performance score is significantly correlated with the mutation rate (**Fig6C&D**), the Spearman’s Rs of 0.53 for EAC and 0.57 for MELA indicates that it is not the only determining factor. We thus interrogated performance for the different mutagenesis clusters for EAC (**FigS16B**). Indeed, the best performance can be observed for the Cluster with mismatch repair deficiencies with a median performance of 36.3. It is followed by Cluster 1, characterized by strong reflux-associated mutagenesis, with a median performance of 27.4. The worst performance is seen for the group with APOBEC-related mutagenesis with a median performance of 10.9. Given the local formation of APOBEC dependent mutation clusters^12,39^, a poor performance was to be expected, since the probability is only to a limited extent dependent on the DNA sequence. Predictions for individual patients enable the assessment of individual loci over the whole cohort (**Fig6E&F**). By assessing the predictions for observed mutations in the main driver mutation locations in EAC and MELA, recurrent mutations are enriched at locations of higher probability of mutagenesis, independent of their location relative to the machine learning sample split (**FigS16D**). At the same time, some mutations are recurrent despite lower mutagenesis probability. Such heterogeneous probabilities across the patient cohort point to individual probabilities of mutagenesis at individual loci as an important factor to be considered when assessing selective pressure.

**Figure 6.**
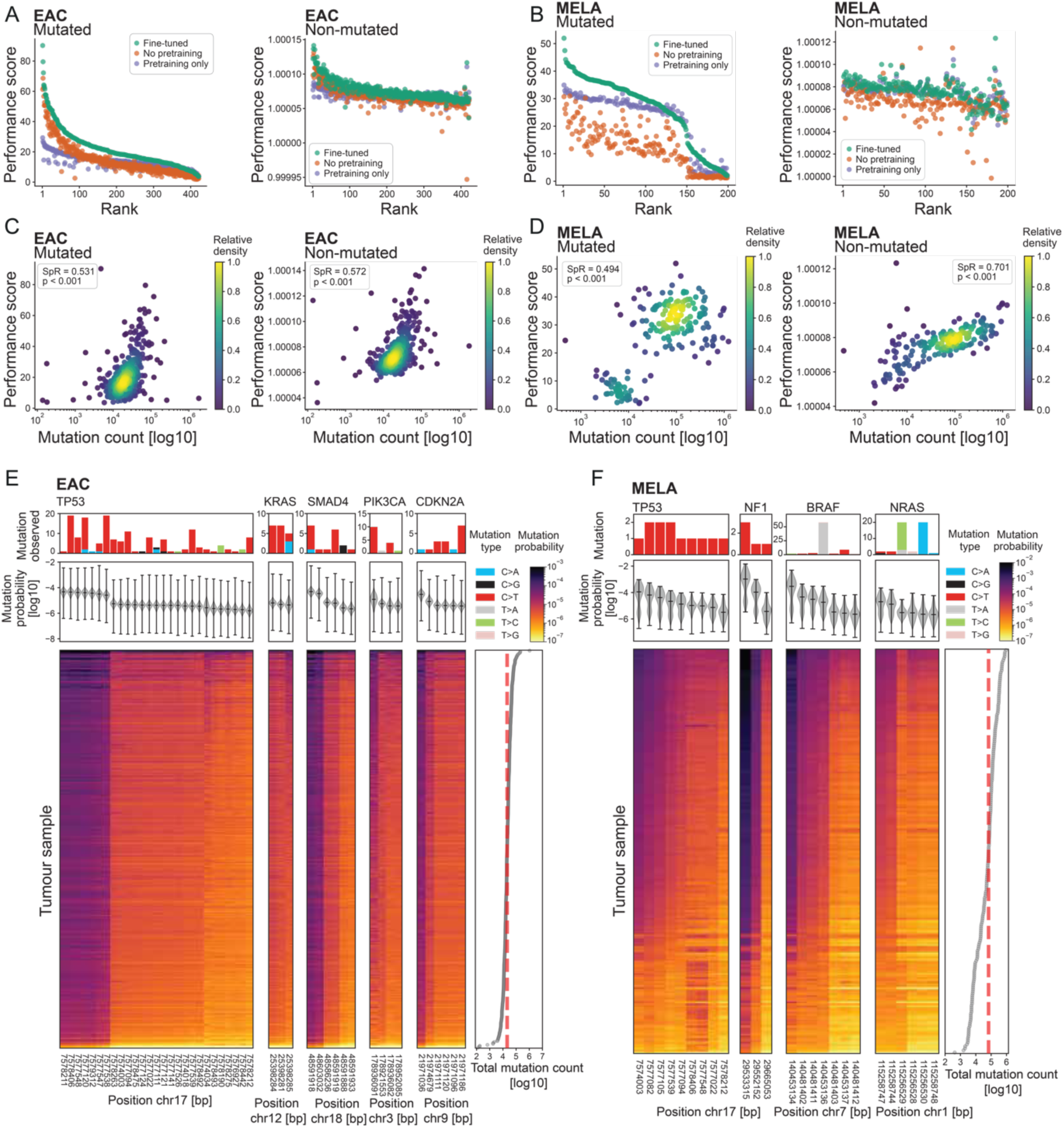
Fine-tuning of EAGLE allows mutation probability assessment on driver mutation sites for individual patients. (A) Performance assessment for individual EAC samples after fine-tuning as the ratio of EAGLE-predicted probability in the test set at sites of observed mutations and sites without observed mutations versus a random model, in comparison to training individual samples without pre-training and applying the pre-trained model to the test set of individual samples without further fine-tuning. (B) Performance assessment for individual MELA samples after fine-tuning analogous to (A). (C) Performance for individual EAC samples in comparison to total SBS counts in the test set at sites of observed mutations and non-mutated sites without observed mutations versus a random model. Sp.R = Spearman’s R. The p-value was derived through a Spearman’s test with an asymptotic t approximation. (D) Performance for individual MELA samples in comparison to total SBS counts analogous to (C). (E) Mutation probabilities for individual EAC patients on sites of *TP53, KRAS, SMAD4, PIK3CA* and *CDKN2A* mutated in the cohort. Observed mutation frequencies are related to the mutation probability over the cohort, with columns ordered by the median probability of the locations, and rows ordered by total mutation count. Only positions classified as known drivers based on the Cancer Genome Interpreter output are shown. (F) Mutation probabilities for individual MELA patients on sites of TP53, NF1, BRAF and NRAS mutated in the cohort. Observed mutation frequencies are related to the mutation probability over the cohort, with columns ordered by the median probability of the locations, and rows ordered by total mutation count. Only positions classified as known drivers based on the Cancer Genome Interpreter output are shown.

Lastly, we investigated whether individual mutation probabilities follow genome biology (**Fig7, FigS17-S23**). Relative to nucleosomes, the EAC samples cluster into groups by the contributions of mutagenesis to either the repetitive mutation probability pattern, or a strong relationship with the linker DNA with distinct differences between non-genic and genic nucleosomes (**Fig7A&B**). For MELA, the C>T rich sample cluster shows a rather homogenous repetitive pattern around nucleosomes for both genic and non-genic nucleosomes. The less mutated MELA sub-group forms a separate cluster with higher mutation rate prediction for genic nucleosome footprints than non-genic (Fig7C&D). Of particular interest is the relative distribution to CTCF binding sites. While we identified coldspots relative to CTCF binding, and with CTCF being a known mutation spot of recurrent mutations^13,14^, analyzing individual samples shows that this is indeed plausible, given that some patients display locally reduced average mutation probabilities and others a local increase both for EAC and MELA (**Fig7E&F**). A similar relationship becomes apparent at TSSs with distinct patterns in patient subgroups that follow mutagenic mechanisms (**Fig7G&H**). Interestingly, this leads to different symmetries of predicted mutation probability relative to the TSS with a higher probability in most EAC and the less heavily mutated MELA subcluster, whereas the heavily mutated MELA samples are reduced in mutagenesis directly after the TSS. Analogous patterns of variable predictions per patient sample can be observed for sequences with the potential to form G4s, TTS, Alu retrotransposons, and exon boundaries (**FigS17&18**). Such individual differences in the relationship with genome biology can be the result of differential exposure to mutagenic mechanisms or individual patient differences, e.g. distinct epigenetic changes or variable cells of origin in a cancer type that can affect the location of mutations even if they originate from the same mechanisms. Inter-sample invariance of mutation patterns from the same mutagenic mechanism can be investigated by focussing on the mutation types with a largely unique etiology. CTT>CGT mutations in esophagus adenocarcinoma are expected to arise as a large majority from acid reflux. Indeed, the distribution relative to genome features reflect largely the data from the EAGLE model on pooled mutations with only limited variance (**FigS19&20**), a homogenous tendency also observed for the largely UV-light-induced TCC>TTC mutations in MELA (**FigS21&22**). There are, however, subtle probability differences which lead to clustering of samples that partially follows mutagenic subgroups. In conclusion, we observe differences in the mutation probability distribution between individual samples that are largely dependent on the mutagenic history of the sample and align with known genome biology.

**Figure 7.**
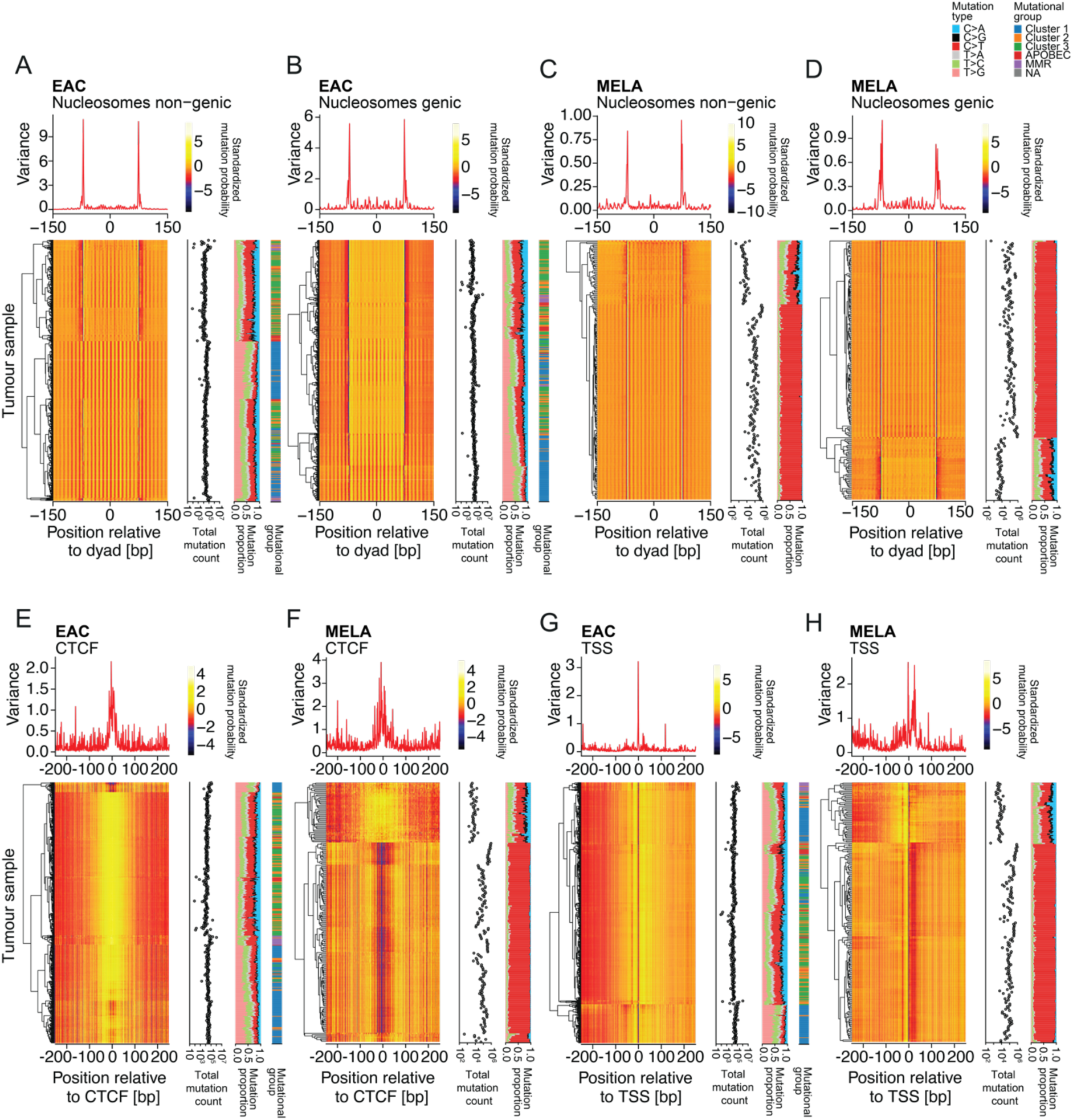
EAGLE predicted mutation probabilities per sample show individual differences of mutagenesis relative to genome-biological features. (A) Predictions of mutation probability relative to non-genic nucleosomes scaled by row. Samples are clustered using hierarchical clustering, set into relation with proportion of mutation types and mutational subgroups. Variance assessed per nucleotide on scaled probabilities. Analogously, data are presented for genic nucleosomes in EAC samples (B), non-genic nucleosomes in MELA samples (C), genic nucleosomes in MELA samples (D), CTCF in EAC samples (E), CTCF in MELA samples (F), TSS in EAC samples (G), and TSS in MELA samples (H). TSS = Transcriptional Start Site, CTCF = CCCTC-Binding Factor.

## Discussion

Understanding the relationship of the genome sequence and its own stability is key to understanding somatic genome evolution. Here, we provide EAGLE, a model able to learn SBS from cancer genomes and infer the probability of mutagenesis for each nucleotide in the genome *en route* to the mutations we observe in individual patient samples. We have found that mutation probabilities, even for the same mutation type in the trinucleotide context, can differ by several orders of magnitude and are the result of sequence motifs for nucleosome positioning, CTCF binding, potential origins of replication, and exon boundaries, as well as direction of transcription and/or replication. The training data was limited to the sequence of the reference genome and incorporated mutations, which allows us to conclude that the footprints of genome biology we observe are the direct or indirect consequence of the sequence itself. For future models, integrating epigenetic data directly into the training may increase performance. However, this design may impact interpretability and introduce bias, because we cannot account for the epigenomes during the whole evolution of the mutation patterns we observe in individual patients. We anticipate that by learning from different modalities, we may differentiate the impact of sequence and epigenetics. Extending EAGLE to additional cancer types and somatic genome evolution in non-cancerous tissues will provide a valuable resource to understand the relationship between sequence and mutagenesis in various contexts. A particular focus will involve studying individual patient samples and germline genome-dependent individual trajectories of somatic genome evolution. In principle, the model can also be used to quantify sequence dependence of artefacts from sequencing and variant-calling, when applied to data of different quality.

SBS arise as a consequence of DNA lesions, the activity of DNA repair mechanisms, and mistakes by polymerases during DNA replication. We demonstrated that sequences which favor nucleosome binding led to variation in mutation rates throughout the different mutation types in EAC and MELA, the extent of which can be individually diverse. The periodicity of mutation probabilities has been shown previously to be the consequence of minor groove orientation relative to nucleosomes, leading to differences in DNA damage impact as well as repair preferences^19^. From a sequence perspective of the same phenomenon, we can first conclude that there is an asymmetry in the sequence motifs underlying nucleosomes, leading to mutation coldspots from various mechanisms, particularly at one of the linkers. One possible explanation for CTT>CGT could be the increased activity of base excision repair at this location to repair the incorporated 8-OxoG^19,37^. However, this explanation would not apply to all mutation types where we observe coldspots at linker sites. Secondly, we can conclude that complex motifs that affect both nucleosome positioning and mutagenesis are distributed over the genome. The locations of such mutation hotspots are typically in intergenic or intronic and late-replicating regions, where functions of strong nucleosome positioning motifs have not been described yet to our knowledge.

Locations experiencing low mutation probability have been challenging to study, because the sparsity of mutation data impeded their identification. They originate from a combination of related sequences linked to genome biology, which is important for gene regulation and chromatin organization. It is therefore not surprising that the genome evolved to keep such sites protected. CTCF binding sites are an interesting case in this context. In individual samples, we predicted them as coldspots as well as sites of locally increased mutation rates, as previously shown^13,14^. There are several possible explanations for this observation. First, the model may get overwhelmed by the large numbers of sites that are unlikely to mutate, which leads to underestimation of the mutagenic binding sites. Also, individual sites may behave differently in the context of mutagenesis, which may become more or less apparent given the different contributions of mutagenic mechanisms in individual patient samples. Second, the observation of increased mutation rates on CTCF sites may not only be a phenomenon of mutagenesis, but also lead to processes of selection, given their potential consequences on gene regulation and instability^13^.

Learning on SBS and 500bp sequence windows, EAGLE is not expected to capture mutation accumulation due to evolutionary selection. Most of the selective pressure has its impact on the protein level, which EAGLE has not been trained to predict. Therefore, the learned probabilities are largely independent of selection and provide a background model of mutagenesis. We can observe that EAGLE predictions of cancer driver genes do not always recapitulate the observed recurrence of these mutations. In the future, it would be interesting to integrate the model output for individual samples as a background for established algorithms to quantify evolutionary selection^40^. The DNA sequence enables epigenetic factors at many levels, leading mutations to specific sequence contexts. However, what lies beyond the sequence, such as epigenetic mechanisms beyond sequence dependence, as well as evolutionary selection, could be used to improve learning.

We provide a framework to learn and understand the DNA sequence foundation for somatic genome evolution, which we aim to develop further towards an integrative understanding of individualized somatic genome evolution and its consequences for somatic mosaicism, age-related diseases, and cancer evolution.

## Methods

Unless otherwise specified as being written in R (v.4.4.1), all code was written in Python (v.3.8.2) with TensorFlow (v.2.5.0), NumPy (v.1.19.2) and Scikit-learn (v.1.0.2)

### Cancer Mutation Data

We used 423 samples from 409 esophageal adenocarcinoma patients and 199 samples from 183 skin melanoma patients from the International Cancer Genome Consortium (ICGC) projects ESAD-UK and MELA-AU, respectively, release 28^29^. SBS mutations for ESAD-UK (n = 12,376,220) and MELA-AU (n = 24,579,703) were mapped relative to the human genome assembly GRCh37 (*hg19*), encompassing the 22 autosomes and both sex chromosomes, unless stated differently.

### Mutational signature calling and mutational subgroup assignment

We decomposed mutational catalogues into 10 distinct SBS mutational signatures using SigProfilerExtractor^30^ and derived relative contributions. Signatures were related to COSMIC signatures (v3.2, https://cancer.sanger.ac.uk/signatures/sbs/). Samples were clustered with k-means clustering (k = 5).

### EAGLE-MUT architecture and training

EAGLE-MUT (<u>E</u>fficient <u>A</u>nalysis with <u>G</u>enome-wide <u>L</u>STM for <u>E</u>valuating <u>MUT</u>ation probabilities, “EAGLE” in the following) is constituted of a bidirectional Long Short-Term Memory (Bi-LSTM) layer^41,42^ with 64 neurons and a 20% dropout rate. It is followed by one fully connected layer with 64 neurons, a customized self-attention layer (https://github.com/CyberZHG/keras-self-attention/tree/master), batch normalization and a ReLU non-linearity layer. The last layer is a fully connected layer with 5 neurons and a softmax activation function.

EAGLE was trained using a categorical cross-entropy loss function with an Adam optimizer, a learning rate of 1e-03, epsilon 1e-07, beta 0.9 on a batch size of 32. EAGLE typically uses a 500bp one-hot encoded DNA reference sequence representing the sequence of As, Cs, Gs, Ts, and Ns, with and without incorporated SBS to predict the observed nucleotide of the somatically evolved genome at every position. The evolved genome was generated by incorporating observed SBS into the reference genome. EAGLE was initially trained on a sample pool combined from the entire cohort. We first divided the whole genome in 500-base-long windows and trained models with 4 random splits of train, validation and test with the proportions 3:1:1. The output is the same 500bp genomic window that was used as input, yet indicating for every position the probabilities of each of the 5 possible nucleotides (including N) after somatic genome evolution.

### EAGLE-MUT genome wide mutation-probability predictions

To calculate mean mutation probabilities for each position in the genome, we used overlapping 500bp windows with a 25bp stride. Probabilities were scaled by the ratio observed/predicted mutations in the training and validation set, and corrected by the proportion of predicted mutations in the whole genome by random sampling to achieve similar patient-specific mutation density. Mutation probabilities were converted to genome browser tracks with the bedGraphToBigWig tool^43^ and visualized with IGV (https://software.broadinstitute.org/software/igv).

### Fine-tuning EAGLE-MUT per sample

For sample-specific assessment, EAGLE was fine-tuned for individual samples. We first selected all 500-base-long windows that contain SBS in individual samples. To create the fine-tuning dataset, the intersection of the splits from the sample pool and the mutated DNA-sequence of individual patient samples was computed. The dataset was complemented with half of the equal amount of SBS-free windows to create balanced splits. Fine-tuning was performed per patient using categorical cross-entropy for 1000 epochs, with early stopping (patience = 10, learning rate = 1e-4, minimum delta = 0). The best-performing weights were restored, which leads to a variable number of training epochs per model.The model of random SBS distribution and models using immediate sequence context.

The model of random SBS distribution assumes no influence of the DNA sequence on the mutation distribution, which leads to a uniform distribution. It was calculated by dividing the number of mutations in the sequence by the sequence length, and the three possible mutation types per nucleotide. To capture the impact of immediate sequence context on mutagenesis, k-mer context models were designed to examine SBS within k-mers of length k=^1,3,5^. These models were calculated by extracting the proportions of the mutation types and dividing them by the frequency of the corresponding context (i.e., the *k-1* adjacent bases, with the mutated base as the central base). The same data split as for EAGLE was used for training, validation and test sets.

### Performance evaluation of mutation probability prediction

We evaluated EAGLE and the *k*-mer-based models by comparing the ratio between the predictions in the test set of each model and the model of random SBS distribution. We predict on 500bp windows, removing 10bp on the edges for comparability of the models and scaling the predictions by the ratio observed/predicted mutations in the training and validation sets. The performance score has the following expression:

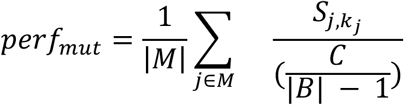

where *M* ⊆ {1, …, *L*} is the set of mutated positions in a sequence of length *L*, and *S*_j,*b*_ is the model output for channel *b* ∈ {*A*, *C*, *G*, *T*, *N*} at position *j*, i.e. the predicted probability of base *b* at that position. *B* = {*A*, *C*, *G*, *T*} is the set of possible destination bases, *k_j_* ∈ *B* is the observed base at mutated position *j*, and *S_j_*_,*k*j_ is the output of the corresponding channel. *C* ∈ [0,1] is the mutation frequency of the random uniform model; under the random model, a position mutates with probability *C* to one of the |*B*| − 1 = 3 alternative bases with equal probability, so *C*/(|*B*| − 1) is the random-model probability of observing base *k_j_*.

Similarly, for positions without a mutation, we quantified prediction performance of positions without an observed mutation using the following expression:

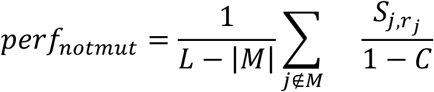

where *r_j_* is the reference base at position *j*, *S_j_*_,*rj*_is the output of the channel corresponding to *r_j_*, and 1 − *C* is the probability under the random model that position *j* remains unmutated.

### Performance assessment with different input window sizes

To investigate the effect of input window size on the performance of EAC EAGLE, we evaluated window sizes from 100 to 1400bp in increments of 100bp. For each window size, the genome was split into training, validation and test sets in 4 random independent iterations, and a model was trained on each split and evaluated on its corresponding test set. Performance was quantified as described above and summarized for each window size as the median across iterations. To visualize the trend across window sizes, a LOESS curve (degree 2, span 0.6) was fitted, and a 95% confidence band was obtained from 1,000 bootstrap refits.

### Performance evaluation with explained variance

The test set was split into batches of 320 bins of 480bp. For each batch, we computed the explained variance between model predictions and observed mutations at single-nucleotide resolution using the explained_variance_score function from scikit-learn. To make values comparable across models, each model’s explained variance was divided by that of a random model, which assigns every non-reference base a uniform mutation rate estimated from the observed mutations in the training and validation windows. Normalized by training plus validation mutation rate, explained variance was computed per batch and averaged across batches.

### Performance evaluation with pseudo R²

Pseudo R² was was computed following the approach of the Roulette model by Seplyarskiy *et al.*^20^, adapting the ‘exp_likelihood_model’, ‘exp_likelihood_null’, ‘avg_likelihood_sampl’ and ‘R²’ functions from the Roulette GitHub repository (https://github.com/vseplyarskiy/Roulette). The EAGLE test set was split into batches of 320 bins, with 480bp per bin. For each position, the predicted mutation probability p was defined as the sum of the probabilities of the three non-reference bases. Within each batch, positions were grouped into 100 quantile bins of p, and the number of sites, the number of observed mutations and the mean p were recorded per bin. Predictions were calibrated by multiplying the mean probability of each bin by a single scaling factor, fitted by maximum binomial likelihood on the corresponding validation batch (binned in the same way) and then applied to the test batch. The null model assigns every position the site-weighted mean calibrated probability. R² was computed as R² = 1 − exp(−2(LL_model − LL_null)), where LL_model is the observed average log-likelihood of the model on the test batch and LL_null is the expected log-likelihood of the null model; R² therefore quantifies the improvement of each model over a uniform mutation rate. R² was computed for all positions, for mutated positions only (log p, with log p̄ as the null reference), and for non-mutated positions only (log(1 − p), with log(1 − p̄) as the null reference). All metrics were computed per batch and averaged across batches; values are shown for each of the four cross-validation splits.

### Area Under the Curve (AUC) and Precision-Recall curve

We used the functions roc_curve and auc from scikit-learn for ROC curves, and precision_recall_curve and average_precision_score for precision-recall curves. In both cases, we subsampled the number of non-mutated sites (n=2,000,000) to match the number of evaluated mutated sites. Thus, our negative samples were the non-mutated sites, and our positive samples were the mutated sites. All probabilities were normalized to the total number of mutations. The same approach was used for the frequency-based *k*-mer models. This performance assessment is designed for classification tasks. It is important to note that the sparsity and stochasticity of mutations impact the use of the derived probabilities as a classifier. This affects the interpretability of absolute AUC values.

### Mutation probability assessment per trinucleotide

To evaluate performance per trinucleotide, we obtained the Spearman correlation between the mean mutation distribution in their trimer context and mean EAGLE predictions. For that, we calculated the mutation type proportions in the test set from EAGLE and the somatically evolved genome.

For EAGLE mutation probability assessment per trinucleotide, predictions for each observed mutation location were aggregated by the underlying trinucleotide and mutation type using R. For assessing relative performance per trinucleotide, we calculated the ratio of probability for correct prediction versus a model assuming random mutation distribution, as described in detail above.

### Mutation assessment relative to cancer driver genes

We selected cancer driver genes that are frequently mutated in esophageal adenocarcinoma (*TP53*, KRAS, *SMAD4*, PIK3CA and *CDKN2A*)^44^ and in melanoma (*TP53*, *NF1*, *BRAF* and *NRAS*), and that also carry mutations in our datasets. We calculated the number of observed mutations per position in these genes, ensuring that each mutation originated from one of 409 and 183 unique donors, in EAC and MELA respectively. For each position associated with an observed mutation, we aggregated the predicted probabilities from all 423 and 199 EAGLE models, respectively, to compare mutation probabilities and recurrence of mutations. We restricted the gene positions to those reported as “known driver” after using the Cancer Genome Interpreter (CGI) pipeline^45^ applied to the SBS mutational data for individual samples with default parameters. To assess whether EAGLE assigns higher probabilities to positions seen during training than to test positions, we used the patient-specific fine-tuned EAGLE models of all 423 EAC patients, trained in four iterations with different genome-wide splits into training, validation and test windows (1,692 models in total). For each observed *TP53* mutation, we extracted the patient’s own predicted probability of the observed mutation type at the mutated nucleotide, in iterations where this position was in the training set and in iterations where it was in the test set. If a position was in the training or test set in more than one iteration, the mean probability was used. Mutations whose position was in the training set in at least one iteration and in the test set in at least one other (n = 121) were compared with a two-sided paired Wilcoxon signed-rank test.

### Genome-wide nucleosome mapping

To assess nucleosomes in genic and non-genic regions we obtained the definitions from Pich et al.^19^ for comparability with their results. However, to cover nucleosome positions for the whole genome equally (chromosomes 1-22), we slightly modified the strategy from Pich *et al.* for genome-wide assessments. In short, we pooled the MNase derived putative nucleosome dyad data from Gaffney *et al*.^46^ using the wigToBigWig and bigWigMerge tools^43^. Then we switched to R (v.4.4.1) and first lifted them over from hg18 to hg19. We smoothed the signal with a 15-bp tri-weight kernel and filtered out low signals with a cut-off of 0.1 and divided by a rolled sum over 151bp to normalize. We then derived the local maxima with a minimal separation of 150bp and reassigned each local maximum to the highest peak in the original data within ±15bp. The final dataset is thus independent of genomic location and is comprised of 9,886,014 nucleosome dyads, which covers 50.4% of the genome’s autosomes, assuming a nucleosome footprint of 147bp and a genome size of hg19 autosomes of 2.9 bio. bp.

### Mutation probability assessment relative to genomic features

Using R (v.4.4.1), mutations were assessed relative to genome biological features, including nucleosomes as defined above. To reduce computing time, a relative subset of 1,000,000 dyads was obtained. CTCF binding sites were obtained as a bed file from https://github.com/bbglab/hotspot_propensity/tree/main/data/ctcf for comparability with Arnedo-Pac *et al*^26^ and lifted over to hg19 using UCSC liftOver^43^. Data were originally obtained from ENCODE^47^ under ID ENCSR000DTR. For melanoma, CTCF binding sites were obtained from ENCODE under ID ENCSR817HTJ (foreskin keratinocytes). Keratinocytes were chosen as the closest skin cell type with high-quality CTCF ChIP-seq data. ETS1 binding sites were obtained from ENCODE under ID ENCSR534VHI (GM23338 induced pluripotent stem cells). As no ETS1 ChIP-seq data are available in ENCODE for melanocytes, this cell line was used as a proxy. Transcriptional Start Sites (TSS), transcription termination sites (TTS), and exon boundaries were derived from TxDb.Hsapiens.UCSC.hg19.knownGene (v.3.2.2) definitions. Alu retrotransposons (Alus) were extracted from the RepeatMasker annotation track, downloaded via the UCSC genome browser (https://www.genome.ucsc.edu/). All Alu families falling between 270 to 330bp in length were retained for analysis. G-quadruplexes (G4s) were defined through sequence prediction using G4Hunter^48^ on autosomes utilizing directly the reference file QP37_hg19_ref.RData provided with the associated R package (n = 343,558). CpG islands were obtained from the UCSC Genome Browser (*hg19*). All CpG positions were split into those inside and outside CpG islands, and the predicted CG>TG probabilities were compared between the two groups with a two-sided Wilcoxon rank-sum test on all CpG positions; violin plots show 100,000 randomly sampled positions per group. Metaprofiles over genome-biological features were generated as the mean coverage of mutations or mutation probabilities relative to the dyads, start sites (TSS, TTS, Alus, exon boundaries) or centers (CTCF binding sites, G4s). For mutation probability assessment using individually fine-tuned models, the analogous analysis was performed in Python.

### Cold- and hotspots definition

Using R, sites of each mutation type in its trinucleotide context were assigned to mutation probability groups based on log10(probability), considering all corresponding trinucleotides on chr1-22 outside RepeatMasker-annotated repeats. Cold- and hotspots were defined as sites below the 1st and above the 99th percentile of the genome-wide distribution, respectively, and the range in between was divided into five groups of equal width on the log10 scale. Genome-wide, EAC CTT>CGT cold- and hotspots each comprise 571,611 sites, corresponding to the 1% of the 57,161,010 million CTT/AAG sites with the lowest and highest predicted probability, respectively; in MELA, TCC>TTC cold- and hotspots each comprise 416,182 of the 41,618,192 million TCC/GGA sites. For comparison, the corresponding trinucleotides with observed mutations were assessed, as well as a random selection of 1,000,000 trinucleotides, irrespective of mutation probability.Assessment of features relative to cold- and hotspots.

Gene element annotation was performed with ChIPSeeker (v1.42.1). Strand annotation relative to transcription was obtained from TxDb.Hsapiens.UCSC.hg19.knownGene (v3.2.2). Replication strand and timing were obtained from OK-Seq data from K562 chronic myeloid leukaemia cells from the reanalysis by Pongor *et al*^49^ downloaded via GEO (GSE131417). Odds ratios were obtained comparing two counts directly or dividing the odds ratio of interest by the odds ratio of the background trinucleotide distribution. P-values were obtained using the Wald test. Coverage of genomic features relative to probability groups was obtained as the mean coverage of nucleosome dyads and exon boundaries as before. CTCF coverage in the healthy gastro-esophageal sphincter was obtained from ENCODE as a pool of ENCFF952NAE and ENCFF629EXD. For melanoma, CTCF coverage in primary foreskin keratinocytes (ENCFF293JXY) and ETS1 coverage in GM23338 skin fibroblast induced pluripotent stem cells (ENCFF731WLT and ENCFF703BCY) were obtained from ENCODE. ORC2 coverage was obtained from Miotto *et al.*^50^, GSE70165 with pooling of the replicates. Further differentiation of defined EAC coldspots was performed by to a TSS within 1kb, binding by CTCF (signal > 9.6) and ORC2 (> 0.85), thresholds corresponding to the 99th percentile of the signal at 1,000,000 random CTT/AAG sites, exon boundaries within 25bp, nucleosome dyads in 73 ± 1bp and 76 ± 1bp. A coldspot can be assigned to more than one group. All coldspots that could not be assigned to a genome-biological feature were assigned to the group “other”. Mutual exclusivity was investigated using overlap between the non-unique coldspot subgroups and assessed for overlaps with UpSetR (v1.4.0). Where not shown separately, coldspots on exon boundaries and linker DNA were pooled within their respective subgroups.

### DNA sequence motif assignments

To derive measures of site-specific sequence content, information content was obtained using seqLogo (v1.72.0). Information content represents the differential between 2 bits and the observed information entropy per nucleotide. Information content was either processed as the sequence logo itself or as aggregated information content of the sequence.

### Transcription factor binding site analyses

CTCF and ETS1 motifs (JASPAR 2024, MA0139.2 and MA0098.4) on chr1-22 were taken from genome-wide predictions for *hg19* overlapping melanocyte DNase I hypersensitive sites (ENCODE ENCSR518JGY). ChIP-seq peaks and signal were obtained from ENCODE for CTCF in primary foreskin keratinocytes (ENCSR817HTJ) and ETS1 in GM23338 induced pluripotent stem cells (ENCSR534VHI). Motifs within a ChIP-seq peak were defined as bound, and motifs without peak overlap and a mean signal ≤ 1.5 (fold change, ± 50 bp) as unbound. Observed mutation rates and EAGLE mutation probabilities were averaged per position relative to the motif centre. Peaks containing a bound motif were centred on the summit and split into mutated (≥ 1 SNV within ± 10 bp) and unmutated peaks. For the heatmaps, peaks were ranked by the mean EAGLE probability within ± 10 bp of the summit and smoothed across neighbouring peaks (Gaussian, σ = 10) for display. Enrichment of mutated peaks along this ranking was tested with fgsea (v1.32.4).

## Code availability

The code and associated tutorials will be made available via Zenodo in full upon publication. It will be shared upon request during the review process.

## Acknowledgements

This work was supported by the Center for Scalable data analytics and artificial intelligence (Scads.AI) Dresden-Leipzig. The authors gratefully acknowledge the computing time made available to them on the high-performance computer at the NHR Center of TU Dresden. This center is jointly supported by the Federal Ministry of Education and Research and the state governments participating in the NHR (www.nhr-verein.de/unsere-partner).

**ARP** and **YDM** were supported by the Mildred Scheel Early Career Center Dresden P2, funded by the German Cancer Aid. **MS** was supported by a TU Dresden Junior Fellowship, a Maria Reiche Postdoctoral fellowship of TU Dresden, and a Marie Skłodowska-Curie Postdoctoral fellowship. **MA** was supported by a DAAD One-Year Grants for Doctoral Candidates.

We thank Jessica Do Amaral Andrade for support on mutational signature calling, the whole Biomedical Genomics group at TU Dresden for fruitful discussions and feedback, as well as Dr. Franziska Baenke for feedback on the manuscript.

## Author contributions

**ARP** has conceptualized the study, **MA**, **YDM**, and **MS** developed the models with input from **ARP**. **MA**, **YDM**, and **ARP** analyzed the data. **ARP** and **MA** wrote the manuscript with input from **MS**.

---

**Figure S1.**
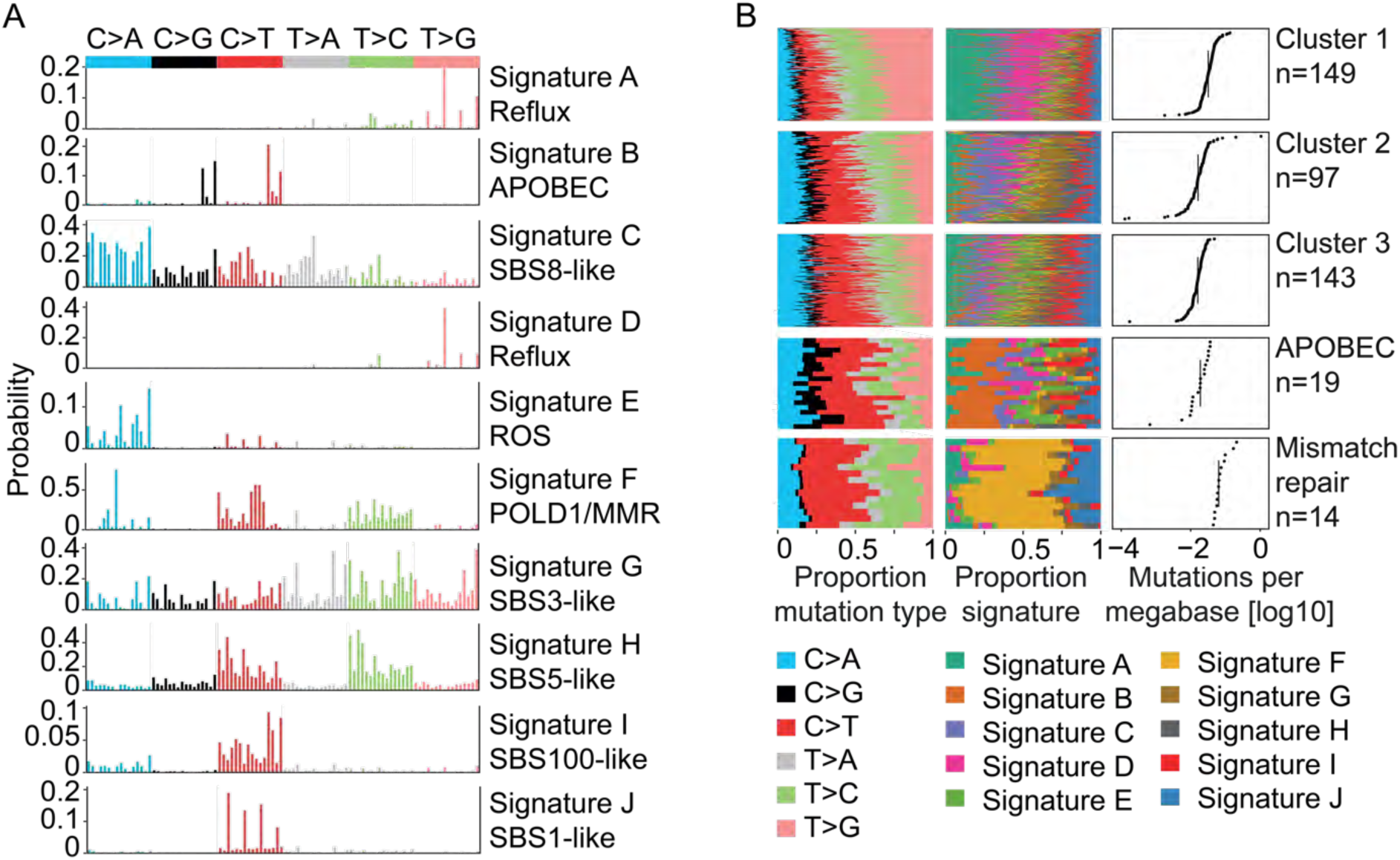
Mutational subgroups in EAC. (A) Mutational signatures present in the cohort. ROS = Reactive Oxygen Species, POLD1/ MMR = Polymerase delta and mismatch repair defects. (B) Proportion of mutation types and signature contributions in the cohort, k-means clustered by the signature contributions and sorted by total mutation rate.

**Figure S2.**
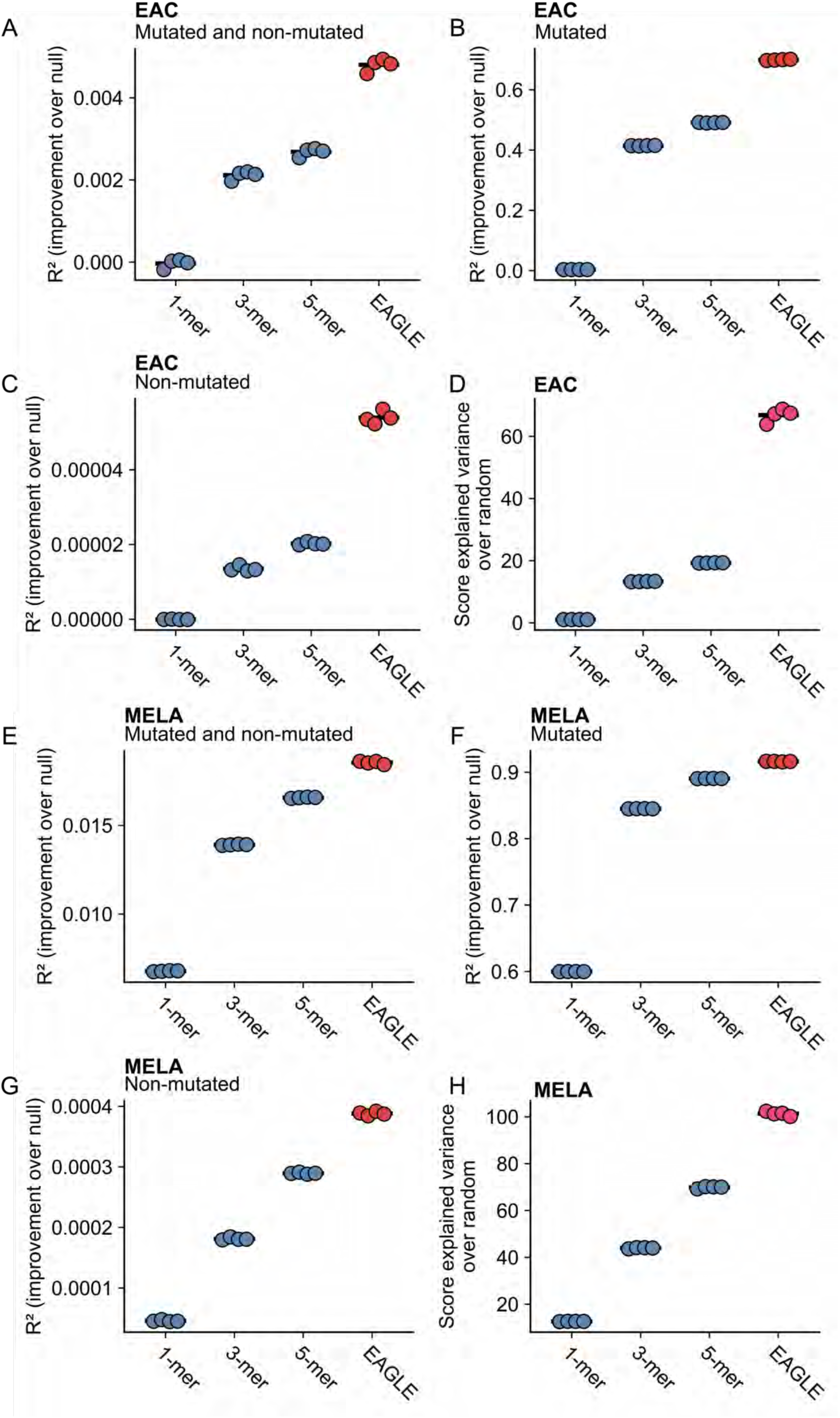
Alternative performance metrics for EAGLE. (A) Performance assessment with Pseudo R^2^ for EAC EAGLE-predicted probability in the test set, aggregated for mutated and non-mutated sites, in comparison to proportional assignment of probabilities based on the observed mutations in the associated k-mer context. (B) Performance assessment with Pseudo R^2^ for EAC EAGLE-predicted probability in the test set at mutated sites, in comparison to proportional assignment of probabilities based on the observed mutations in the associated k-mer context. (C) Performance assessment with Pseudo R^2^ for EAC EAGLE predictions in the test set at non-mutated sites, in comparison to proportional assignment of probabilities based on the observed mutations in the associated k-mer context. (D) Performance assessment using a score for explained variance in the test set relative to EAC EAGLE predictions based on proportional mutation distribution in a random model, in comparison to proportional assignment of probabilities based on the observed mutations in the associated k-mer context. (E) Performance assessment with Pseudo R^2^ for MELA EAGLE-predicted probability in the test set, aggregated for mutated and non-mutated sites, in comparison to proportional assignment of probabilities based on the observed mutations in the associated k-mer context. (F) Performance assessment with Pseudo R^2^ for MELA EAGLE-predicted probability in the test set at mutated sites, in comparison to proportional assignment of probabilities based on the observed mutations in the associated k-mer context. (G) Performance assessment with Pseudo R^2^ for MELA EAGLE predictions in the test set at non-mutated sites, in comparison to proportional assignment of probabilities based on the observed mutations in the associated k-mer context (H) Performance assessment using a score for explained variance in the test set relative to MELA EAGLE predictions based on proportional mutation distribution in a random model, in comparison to proportional assignment of probabilities based on the observed mutations in the associated k-mer context.

**Figure S3.**
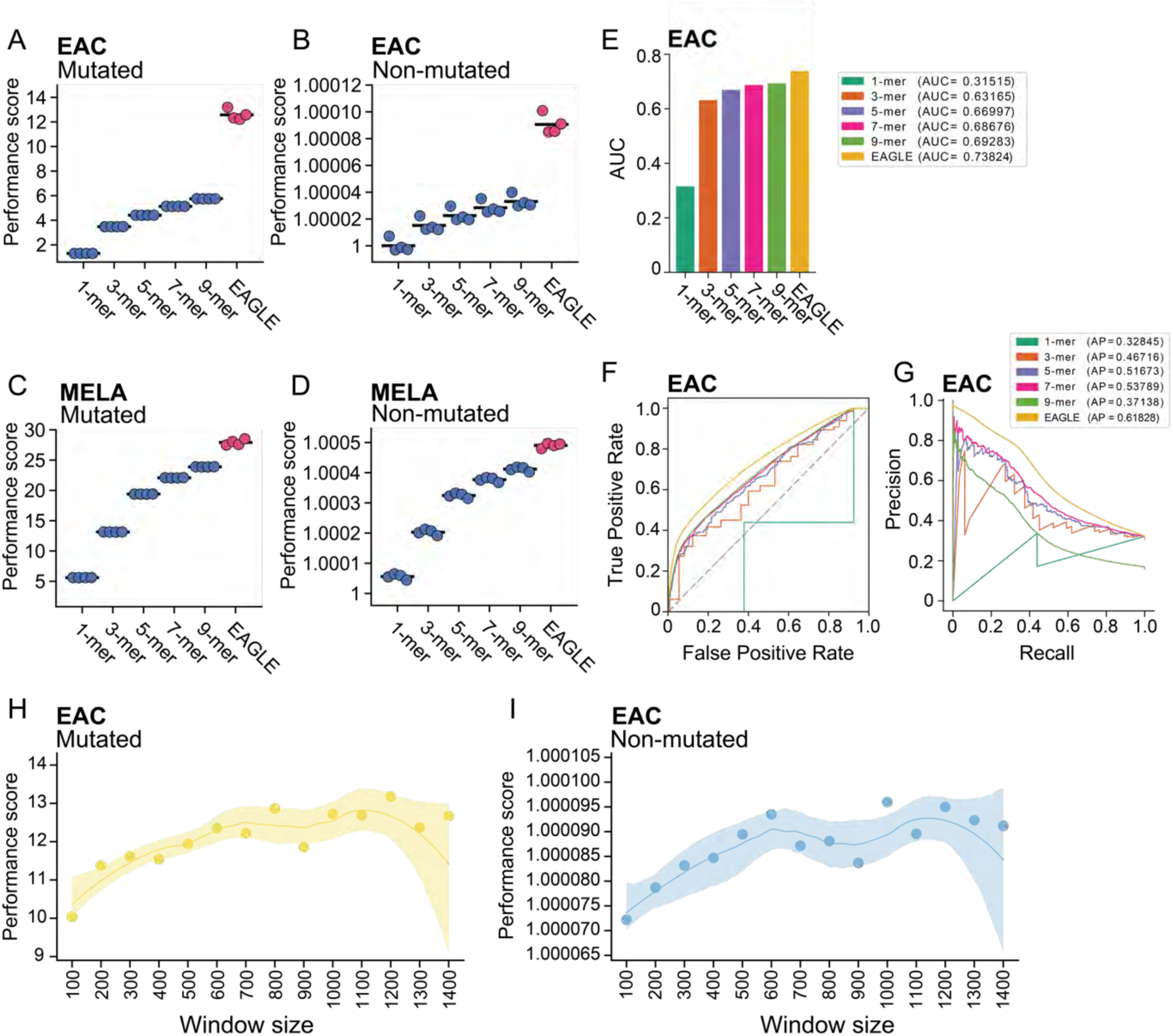
EAGLE assessment relative to context range. (A) Performance assessment as the ratio of EAGLE-predicted probability in the test set at mutated sites in EAC versus a random model, in comparison to proportional assignment of probabilities based on the observed mutations in the associated k-mer context, including 7-mer and 9-mer. 4 models are shown, trained with 4 different splits of training, validation, and test sets. (B) Performance assessment of non-mutated sites in EAC, analogous to (A). (C) Performance assessment as the ratio of EAGLE-predicted probability in the test set at mutated sites in MELA versus a random model, in comparison to proportional assignment of probabilities based on the observed mutations in the associated k-mer context, including 7-mer and 9-mer. 4 models are shown, trained with 4 different splits of training, validation, and test sets. (D) Performance assessment of non-mutated sites in MELA, analogous to (C). (E) Area under the ROC curve (AUC) of EAGLE and the frequency-based k-mer models in predicting mutated versus non-mutated sites in the EAC test set. (F) Receiver operating characteristic curve (ROC) for the models in (E). The dashed diagonal line indicates the performance of a random classifier. (G) Precision-recall curves for the models in (E). Average precision (AP) for each model is indicated in the legend. (H) Performance assessment as the ratio of EAGLE-predicted probability in the test set at mutated sites in EAC versus a random model, in comparison to different input window sizes for EAGLE. Depicted with solid points is mean performance across 4 different splits of training, validation, and test sets; with a solid line depicting a loess fit and the 95% confidence interval as shading. (I) Performance assessment of non-mutated sites in EAC, analogous to (H).

**Figure S4.**
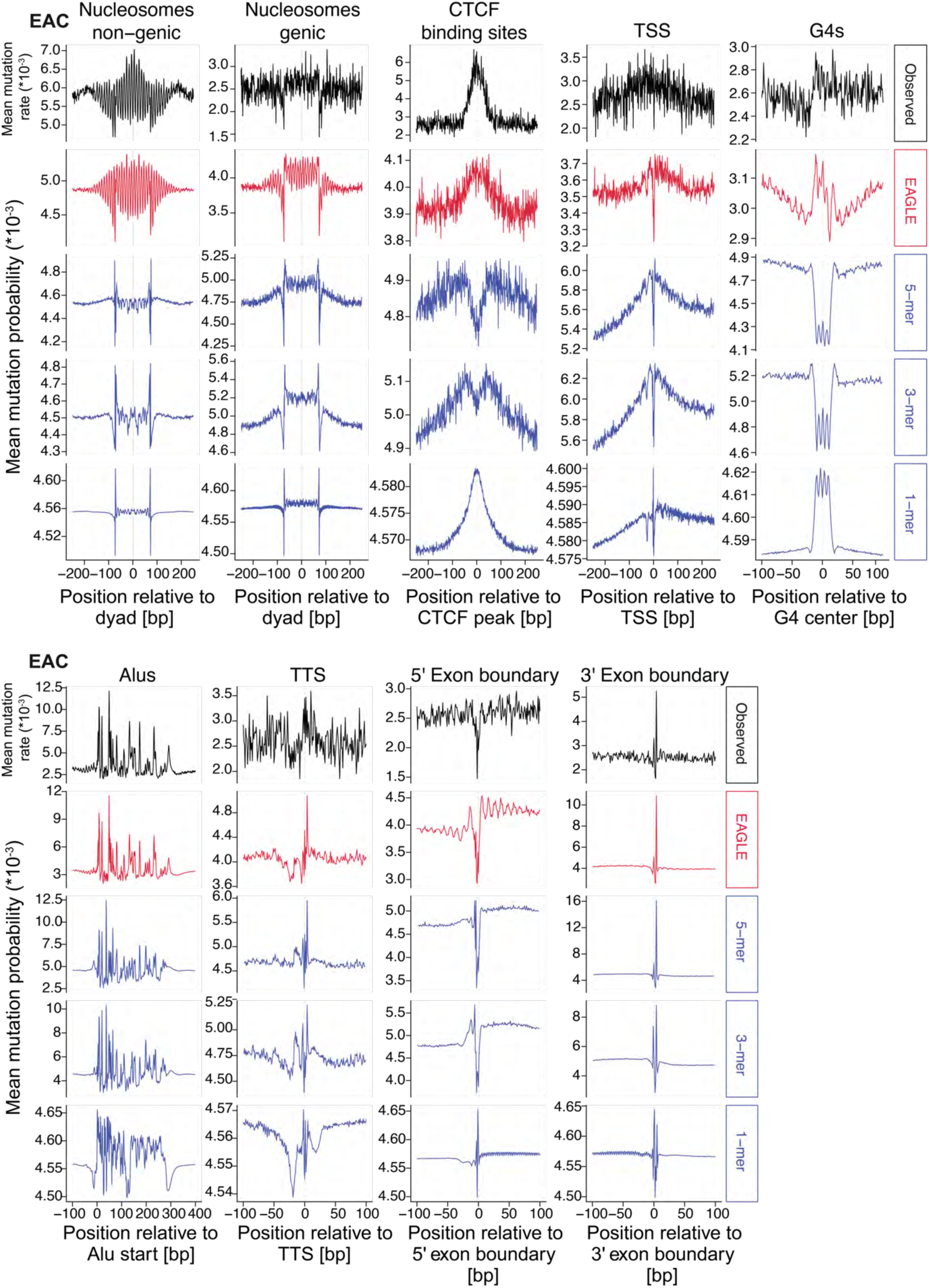
EAGLE predictions for EAC relative to genome elements. Mean mutation rate distribution of EAGLE predictions in comparison to mean observed SBS distribution and proportional assignment of probabilities based on the observed mutations in the associated k-mer context, relative to genome elements. CTCF = CCCTC-Binding Factor, TSS = Transcriptional Start Site, G4s = predicted G-quadruplex structures, Alus = Alu retrotransposons, TTS = Transcriptional Termination Site.

**Figure S5.**
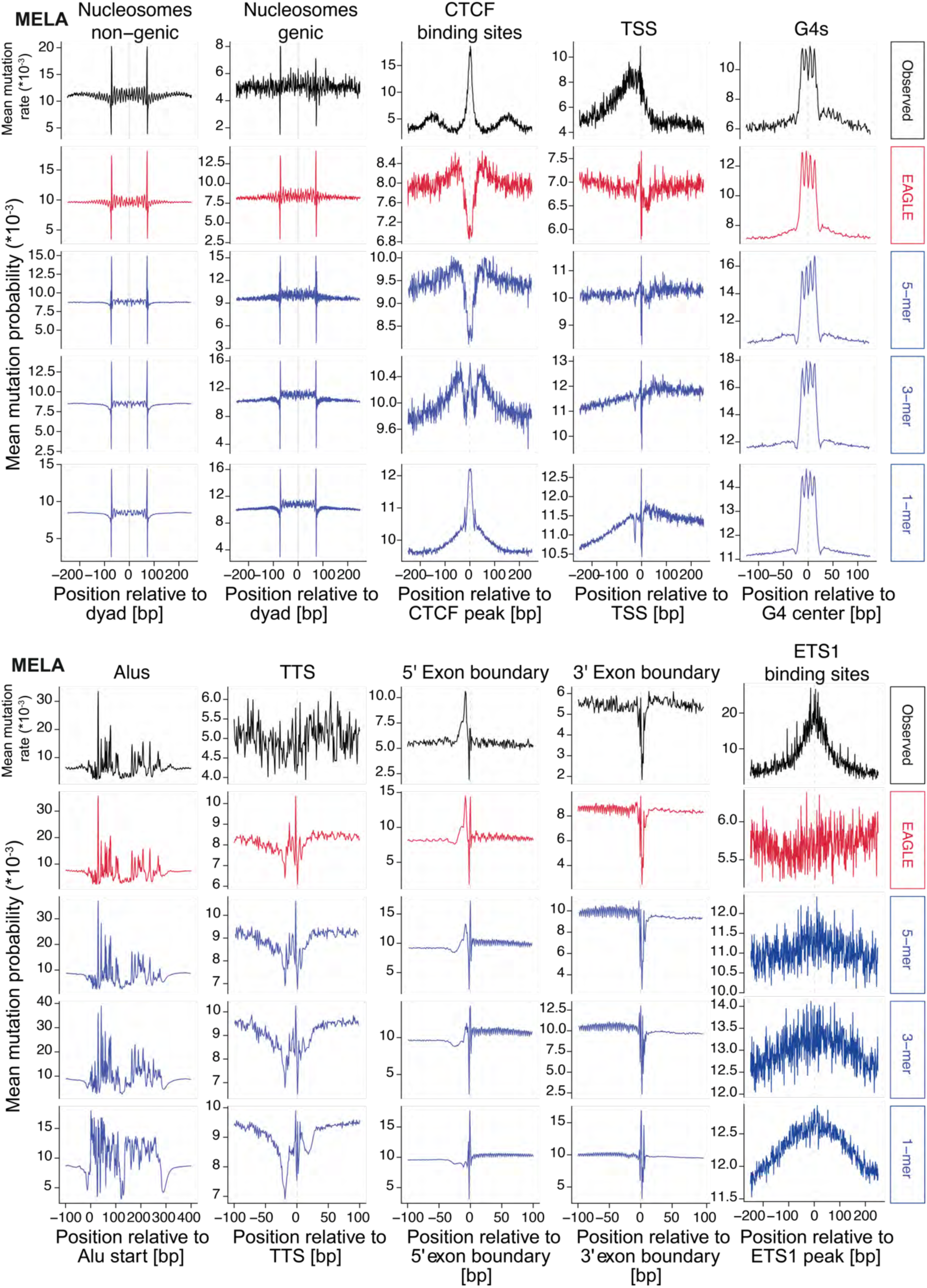
EAGLE predictions for MELA relative to genome elements. Mean mutation rate distribution of EAGLE predictions in comparison to mean observed SBS distribution and proportional assignment of probabilities based on the observed mutations in the associated k-mer context, relative to genome elements. CTCF = CCCTC-Binding Factor, TSS = Transcriptional Start Site, G4s = predicted G-quadruplex structures, Alus = Alu retrotransposons, TTS = Transcriptional Termination Site, ETS1 = ETS1 Binding Factor.

**Figure S6.**
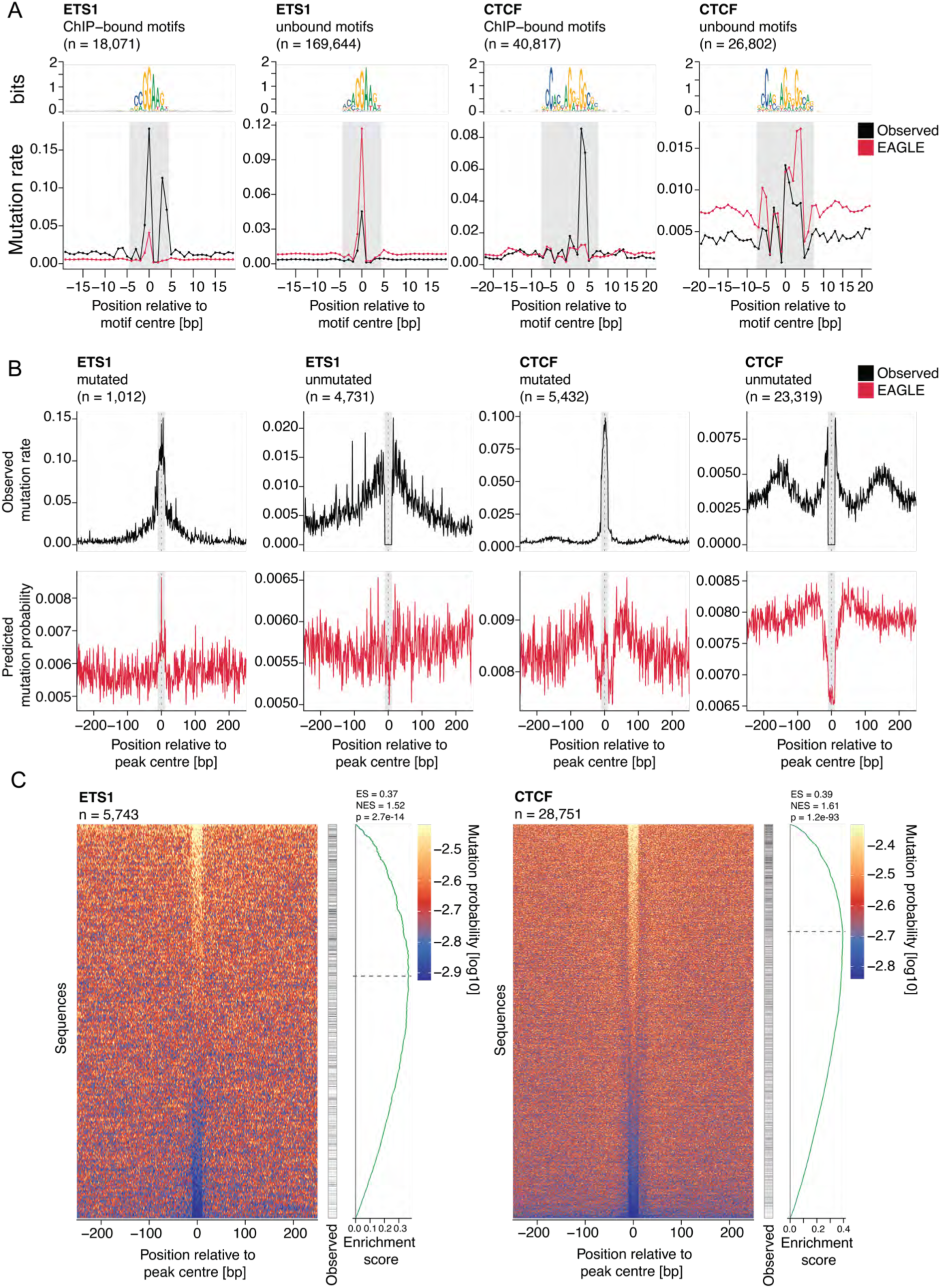
EAGLE prediction assessment for ETS1 and CTCF binding sites in MELA. (A) Mean mutation rate distribution of EAGLE predictions in comparison to mean observed SBS distribution, relative to the center of ChIP-seq-bound and unbound ETS1 and CTCF logos show the motif sequence; n, number of motif sites. (B) Mean mutation rate distribution of EAGLE predictions (bottom) in comparison to mean observed SBS distribution (top), relative to the summit of ChIP-seq peaks containing bound ETS1 or CTCF motif, separated into mutated (≥ 1 SBS within ± 10bp of the summit) and unmutated peaks; n, number of peaks. (C) EAGLE-predicted mutation probabilities in log10 scale for all ChIP-seq peaks in (B) ranked by the prediction within ± 10bp of the summit. Right: mutated peaks in black, and their running enrichment score along the ranking in green. ES = enrichment score, NES = normalized enrichment score, p-values were obtained by permutation of random peak sets of the same size.

**Figure S7.**
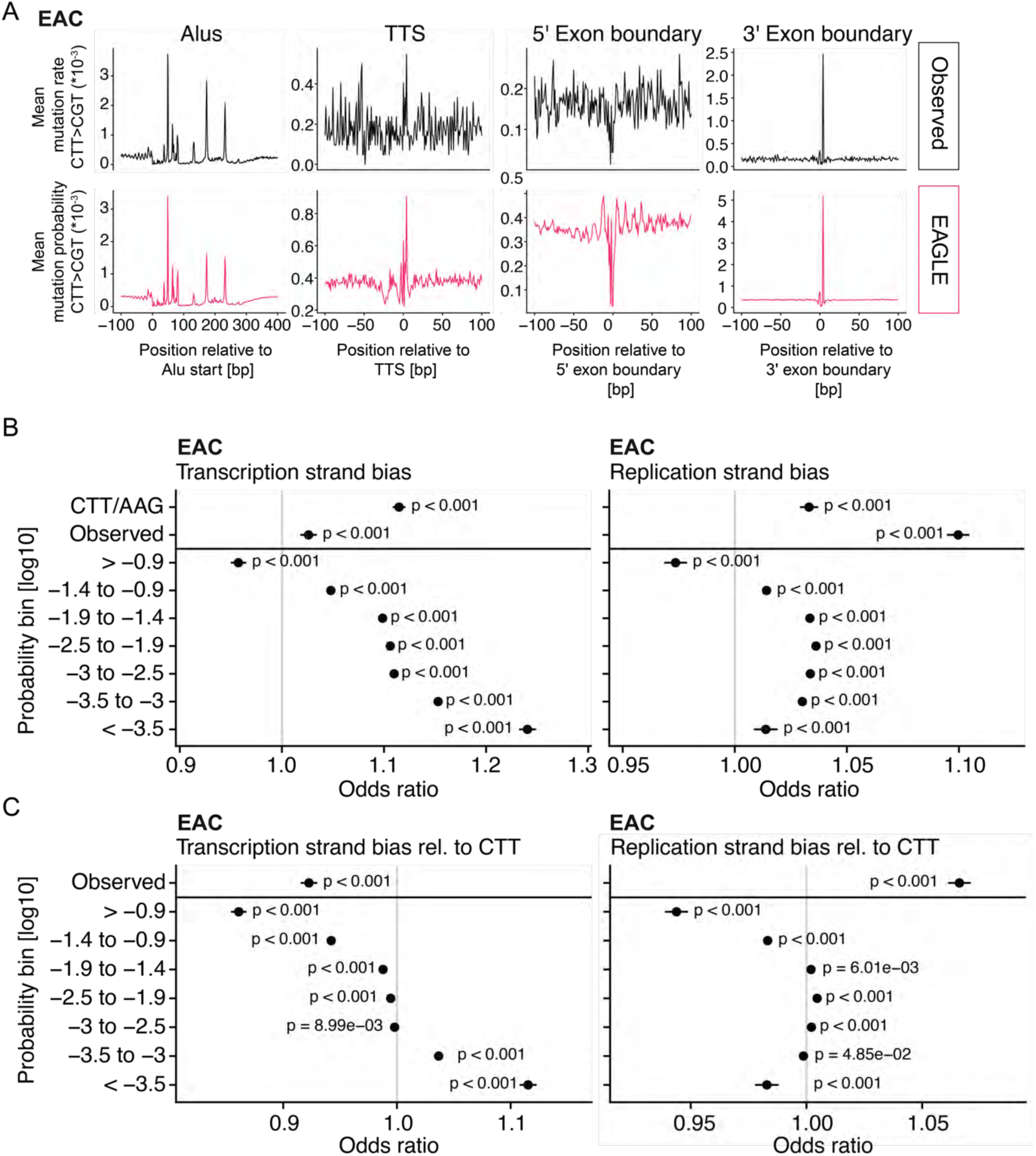
CTT>CGT mutation probabilities in EAC follow distinct sequence patterns and strand biases. (A) Mean mutation rate distribution of EAGLE predictions for CTT>CGT/AAG>ACG in comparison to mean observed SBS distribution, relative to genome elements. Alus = Alu retrotransposons, TTS = Transcriptional Termination Site. (B) Transcription- and DNA replication strand bias of EAGLE predictions for CTT>CGT/AAG>ACG in comparison to mean observed SBS distribution and background distribution of CTT/AAG. Odds ratios are calculated with associated p-values based on the imbalance of the strands. (C) Transcription- and DNA replication strand bias of EAGLE predictions for CTT>CGT/AAG>ACG in comparison to mean observed SBS distribution. Odds ratios are calculated with associated p-values based on the imbalance of the strands, corrected for the imbalance of the underlying trinucleotide.

**Figure S8.**
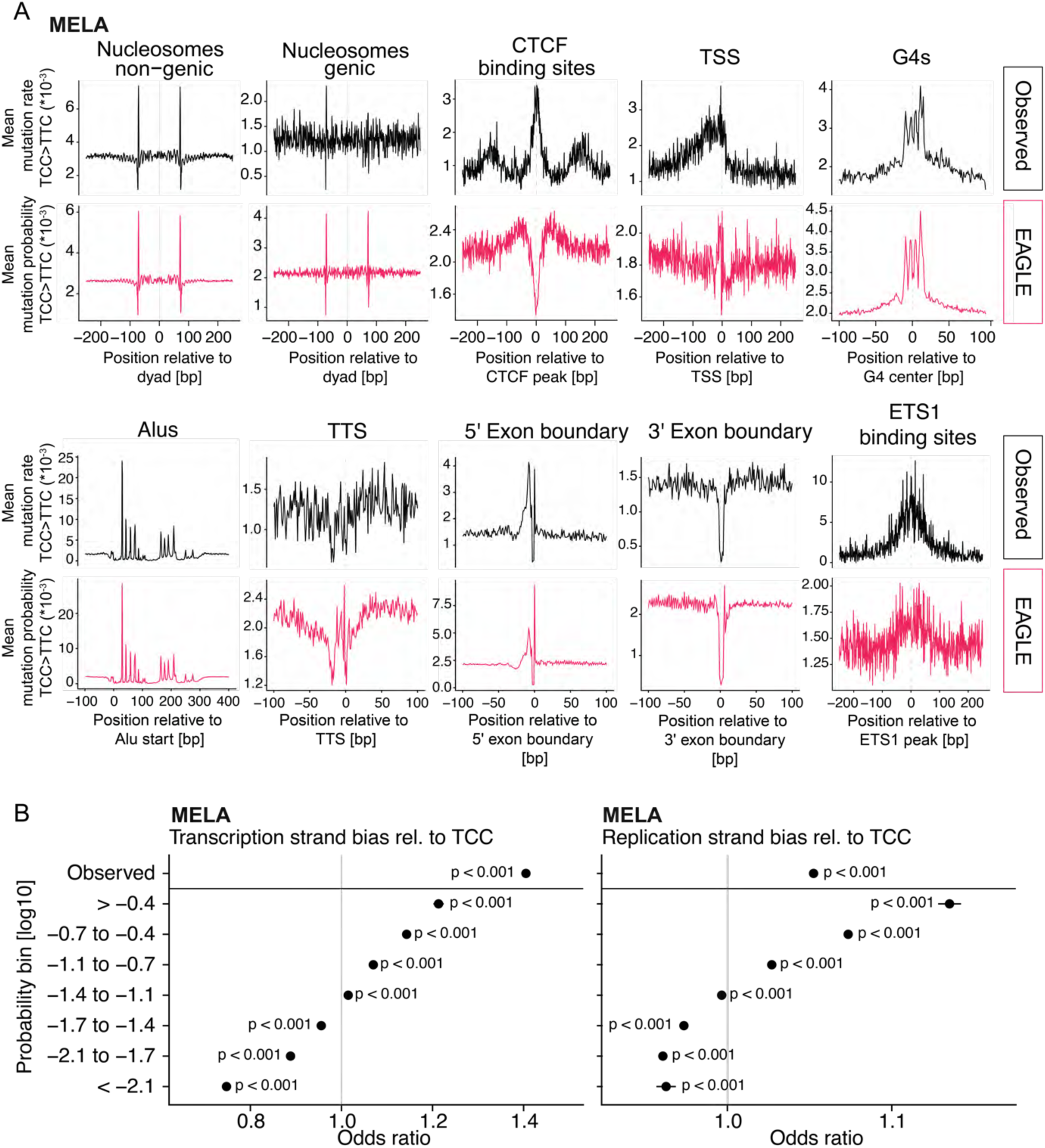
TCC>TTC mutation probabilities in MELA follow distinct sequence patterns and strand biases. (A) Mean mutation rate distribution of EAGLE predictions for TCC>TTC/GGA>GAA in comparison to mean observed SBS distribution, relative to genome elements. CTCF = CCCTC-Binding Factor, TSS = Transcriptional Start Site, G4s = predicted G-quadruplex structures, Alus = Alu retrotransposons, TTS = Transcriptional Termination Site, ETS1 = ETS1 Binding Factor. (B) Transcription- and DNA replication strand bias of EAGLE predictions for TCC>TTC/GGA>GAAin comparison to mean observed SBS distribution. Odds ratios are calculated with associated p-values based on the imbalance of the strands, corrected for the imbalance of the underlying trinucleotide.

**Figure S9.**
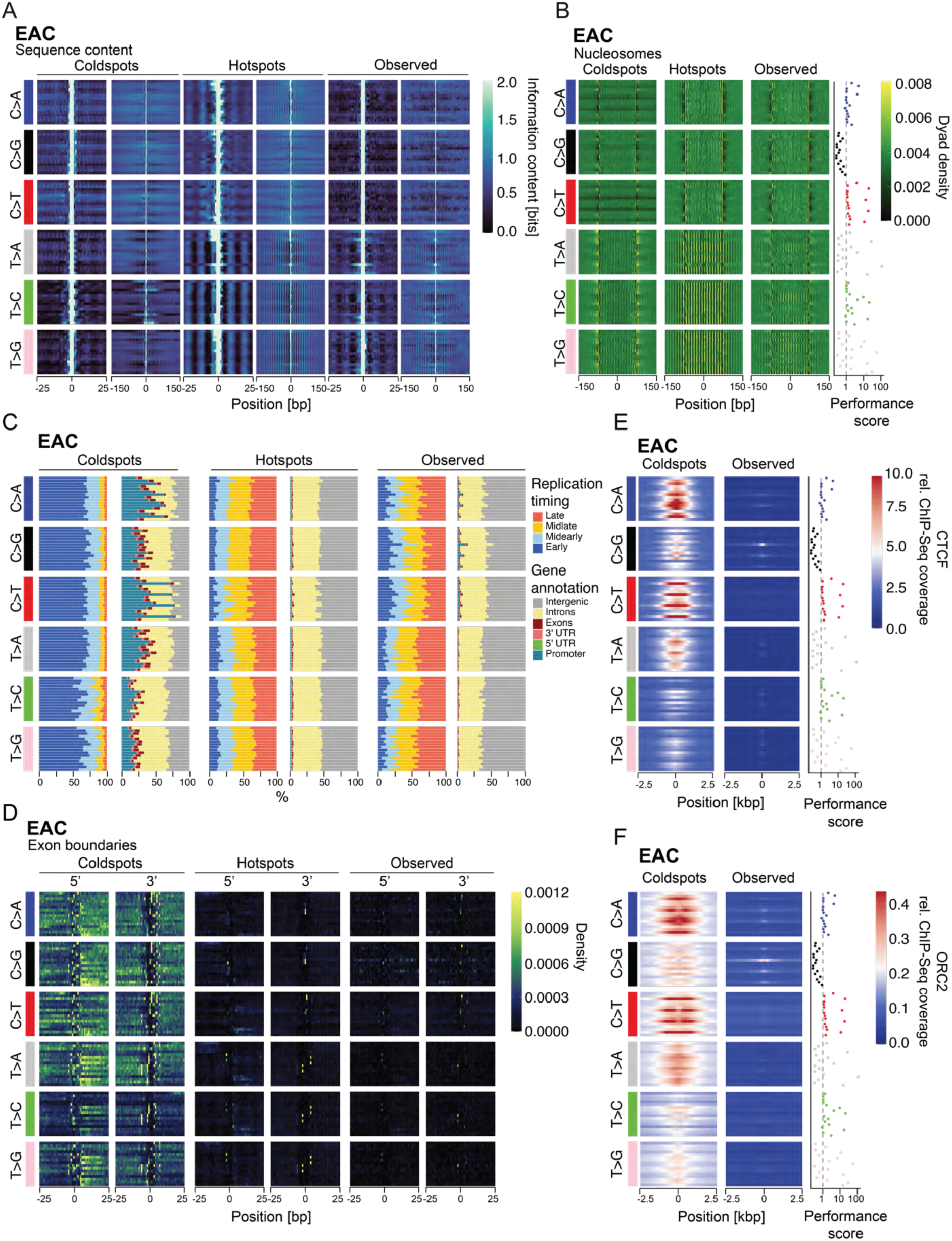
Mutation cold- and hotspots in EAC follow distinct sequence patterns related to genome biology. (A) Sequence motif patterns for coldspots, hotspots and observed SBS, differentiated by their mutation type and trinucleotide context as before. Information content represents the differential of 2 bits and the information entropy of the sequences flanking the mutated trinucleotide (B) Nucleosome dyad density for coldspots, hotspots and observed SBS, differentiated by their mutation type and trinucleotide context as before. (C) Gene- and DNA replication timing-dependent annotation of coldspots, hotspots and observed SBS, differentiated by their mutation type and trinucleotide context as before. (D) Density of exon boundaries at coldspots, hotspots and observed SBS, differentiated by their mutation type and trinucleotide context as before. (E) Relative ChiP-Seq coverage of CTCF at coldspots, and observed SBS, differentiated by their mutation type and trinucleotide context as before. Hotspots are not shown due to no coverage. CTCF = CCCTC-Binding Factor. (F) Relative ChiP-Seq coverage of ORC2 at coldspots, and observed SBS, differentiated by their mutation type and trinucleotide context as before. ORC2 represents potential origins of DNA replication. Hotspots are not shown due to no coverage. Performance as the ratio of EAGLE versus a random model for each trinucleotide is shown per row. The dashed gray line at 1 indicates equal performance to the random model.

**Figure S10.**
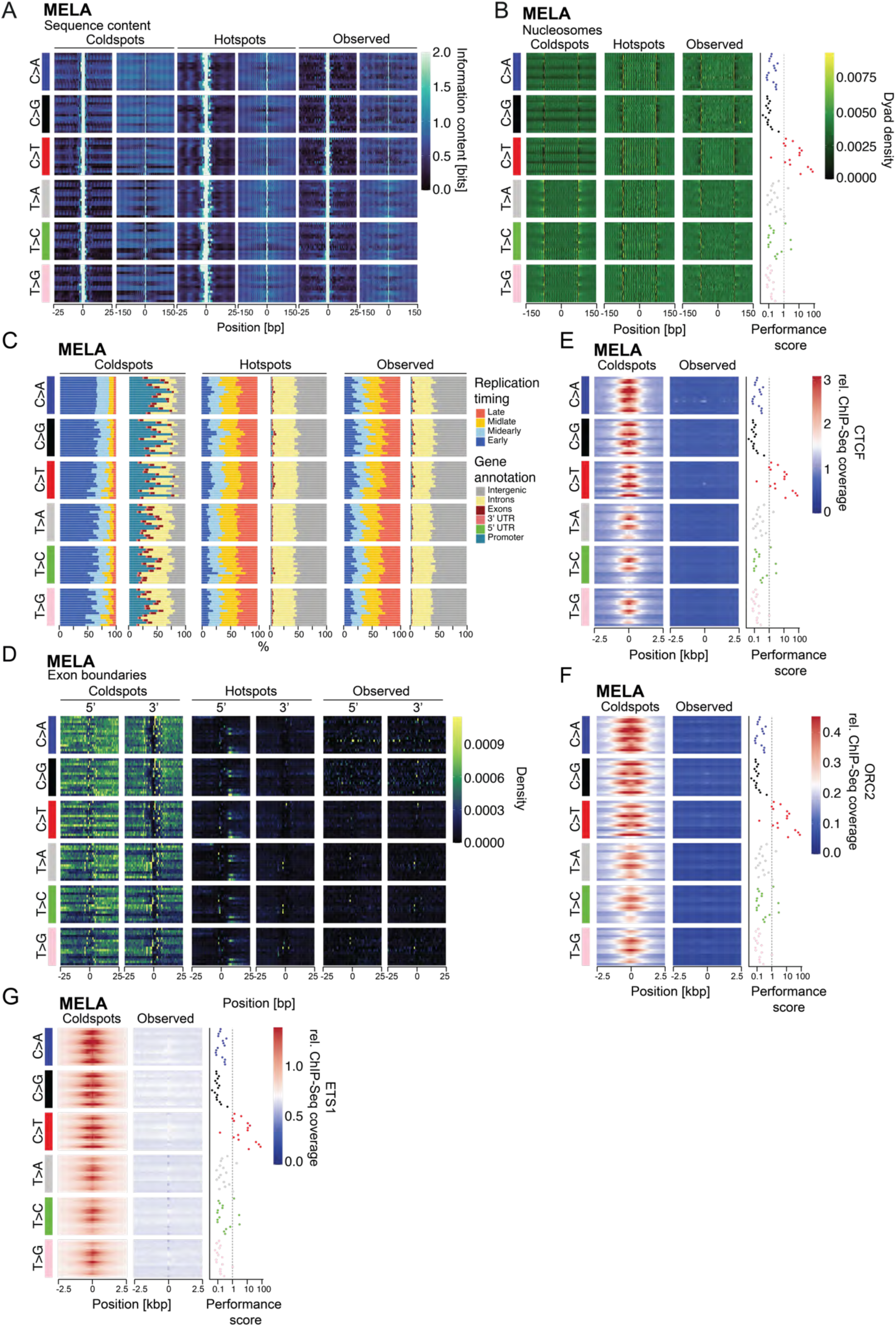
Mutation cold- and hotspots in MELA follow distinct sequence patterns related to genome biology. (A) Sequence motif patterns for coldspots, hotspots and observed SBS, differentiated by their mutation type and trinucleotide context as before. Information content represents the differential of 2 bits and the information entropy of the sequences flanking the mutated trinucleotide (B) Nucleosome dyad density for coldspots, hotspots and observed SBS, differentiated by their mutation type and trinucleotide context as before. (C) Gene- and DNA replication timing-dependent annotation of coldspots, hotspots and observed SBS, differentiated by their mutation type and trinucleotide context as before. (D) Density of exon boundaries at coldspots, hotspots and observed SBS, differentiated by their mutation type and trinucleotide context as before. (E) Relative ChiP-Seq coverage of CTCF at coldspots, and observed SBS, differentiated by their mutation type and trinucleotide context as before. Hotspots are not shown due to no coverage. CTCF = CCCTC-Binding Factor. (F) Relative ChiP-Seq coverage of ORC2 at coldspots, and observed SBS, differentiated by their mutation type and trinucleotide context as before. ORC2 represents potential origins of DNA replication. Hotspots are not shown due to no coverage. (G) Relative ChiP-Seq coverage of ETS1 at coldspots, and observed SBS, differentiated by their mutation type and trinucleotide context as before. Hotspots are not shown due to no coverage. Performance as the ratio of EAGLE versus a random model for each trinucleotide is shown per row. The dashed gray line at 1 indicates equal performance to the random model.

**Figure S11.**
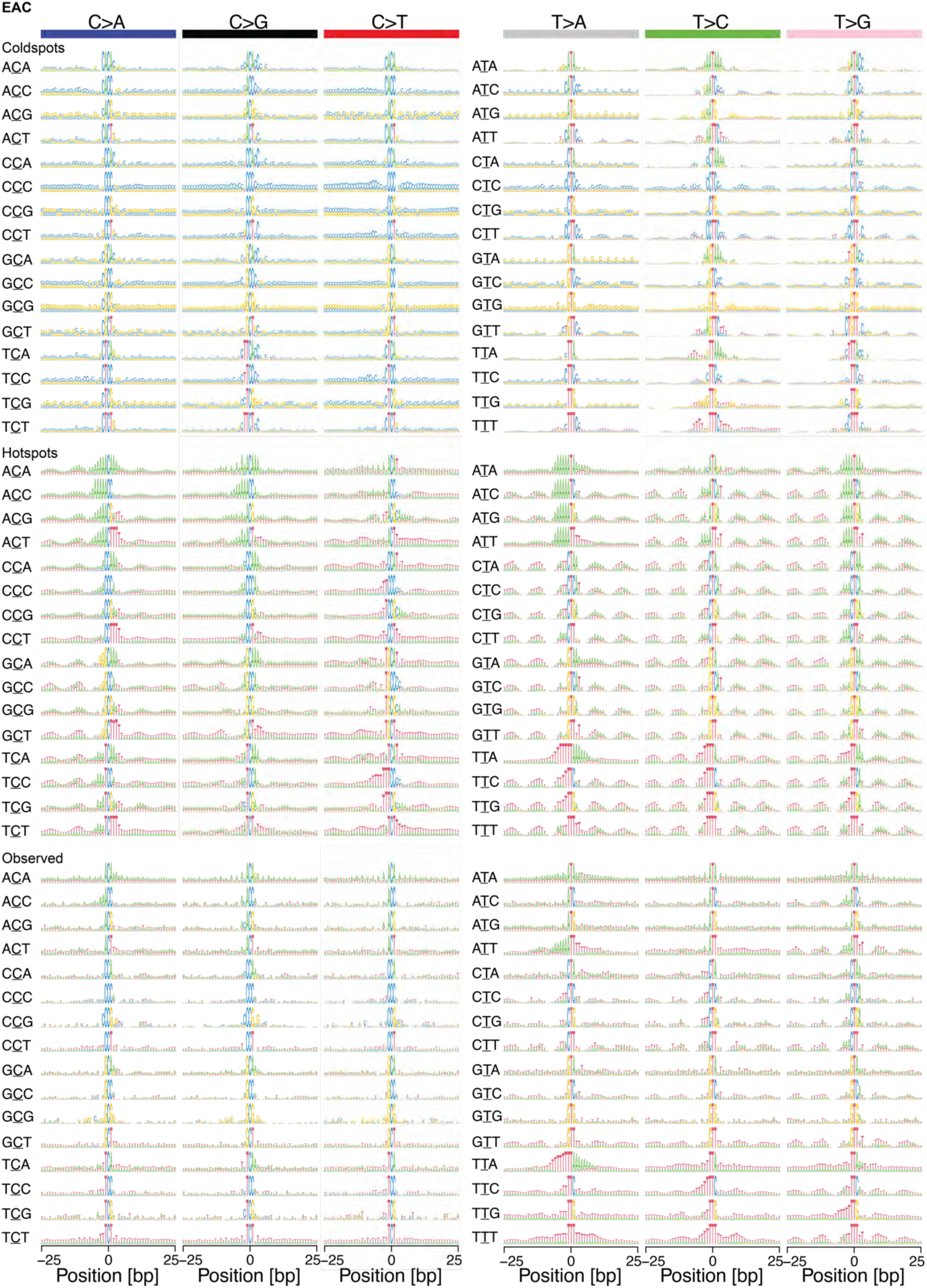
Mutation cold- and hotspots in EAC follow distinct immediate sequence motifs. Sequence motif patterns of 50bp for coldspots, hotspots and observed SBS, differentiated by their mutation type and trinucleotide context as before. Information content represents the differential of 2 bits and the information entropy of the sequences flanking the mutated trinucleotide, resolved by nucleotide as a motif visualization. The data correspond to the aggregated information content of FigureS9.

**Figure S12.**
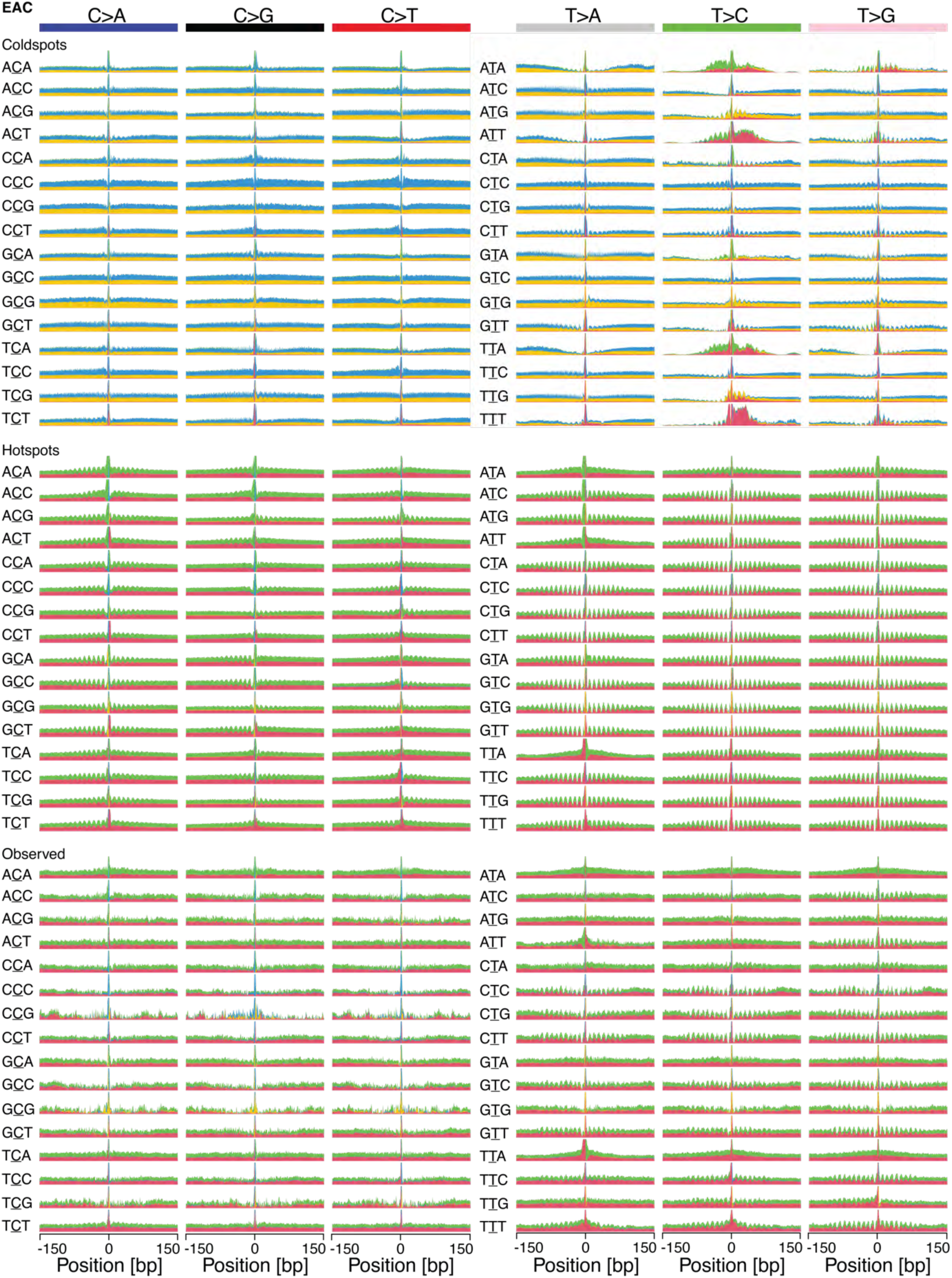
Mutation cold- and hotspots in EAC follow distinct intermediate range sequence motifs. Sequence motif patterns of 300bp for coldspots, hotspots and observed SBS, differentiated by their mutation type and trinucleotide context as before. Information content represents the differential of 2 bits and the information entropy of the sequences flanking the mutated trinucleotide, resolved by nucleotide as a motif visualization. The data correspond to the aggregated information content of FigureS9.

**Figure S13.**
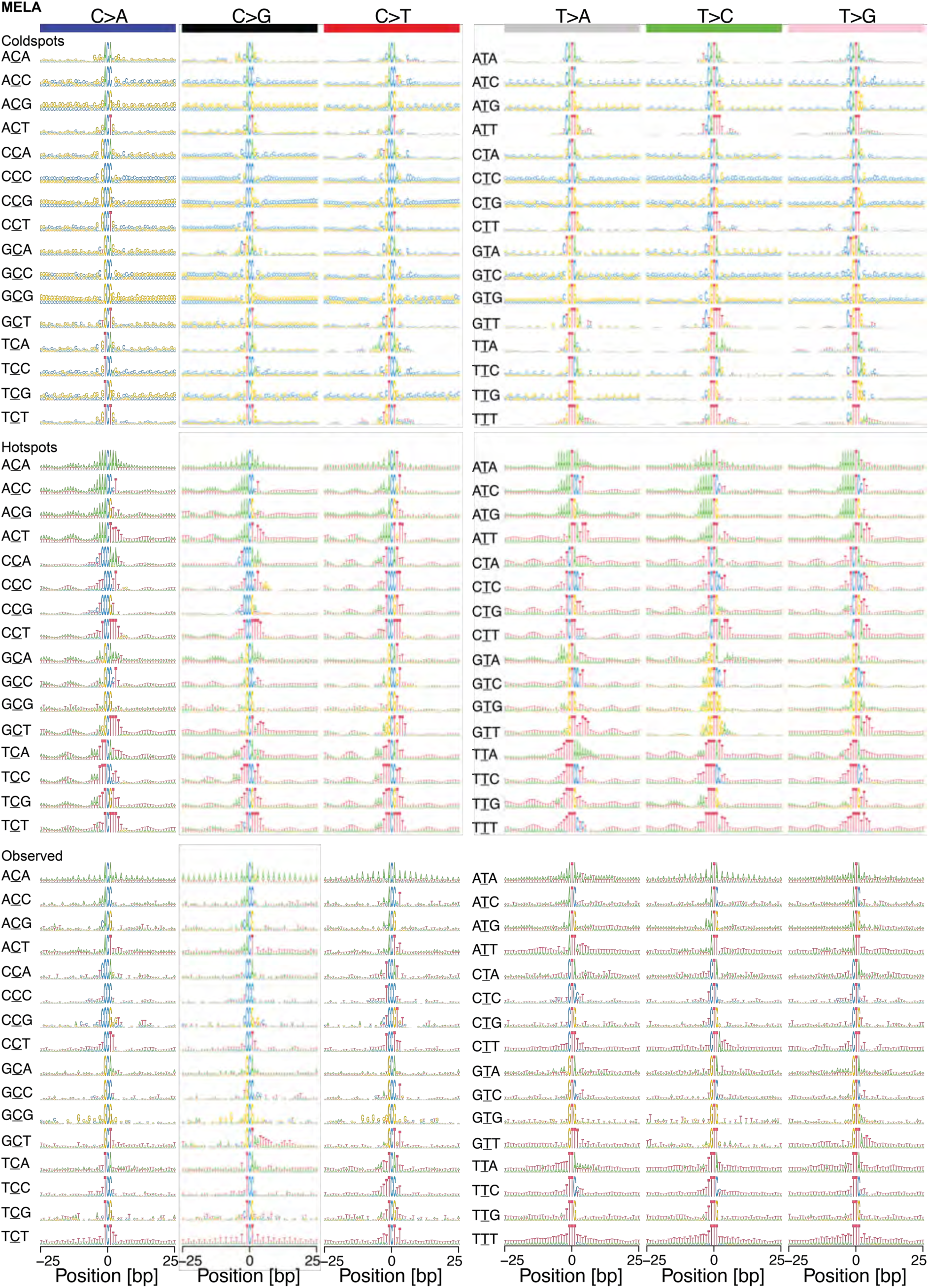
Mutation cold- and hotspots in MELA follow distinct immediate sequence motifs. Sequence motif patterns of 50bp for coldspots, hotspots and observed SBS, differentiated by their mutation type and trinucleotide context as before. Information content represents the differential of 2 bits and the information entropy of the sequences flanking the mutated trinucleotide, resolved by nucleotide as a motif visualization. The data correspond to the aggregated information content of FigureS10.

**Figure S14.**
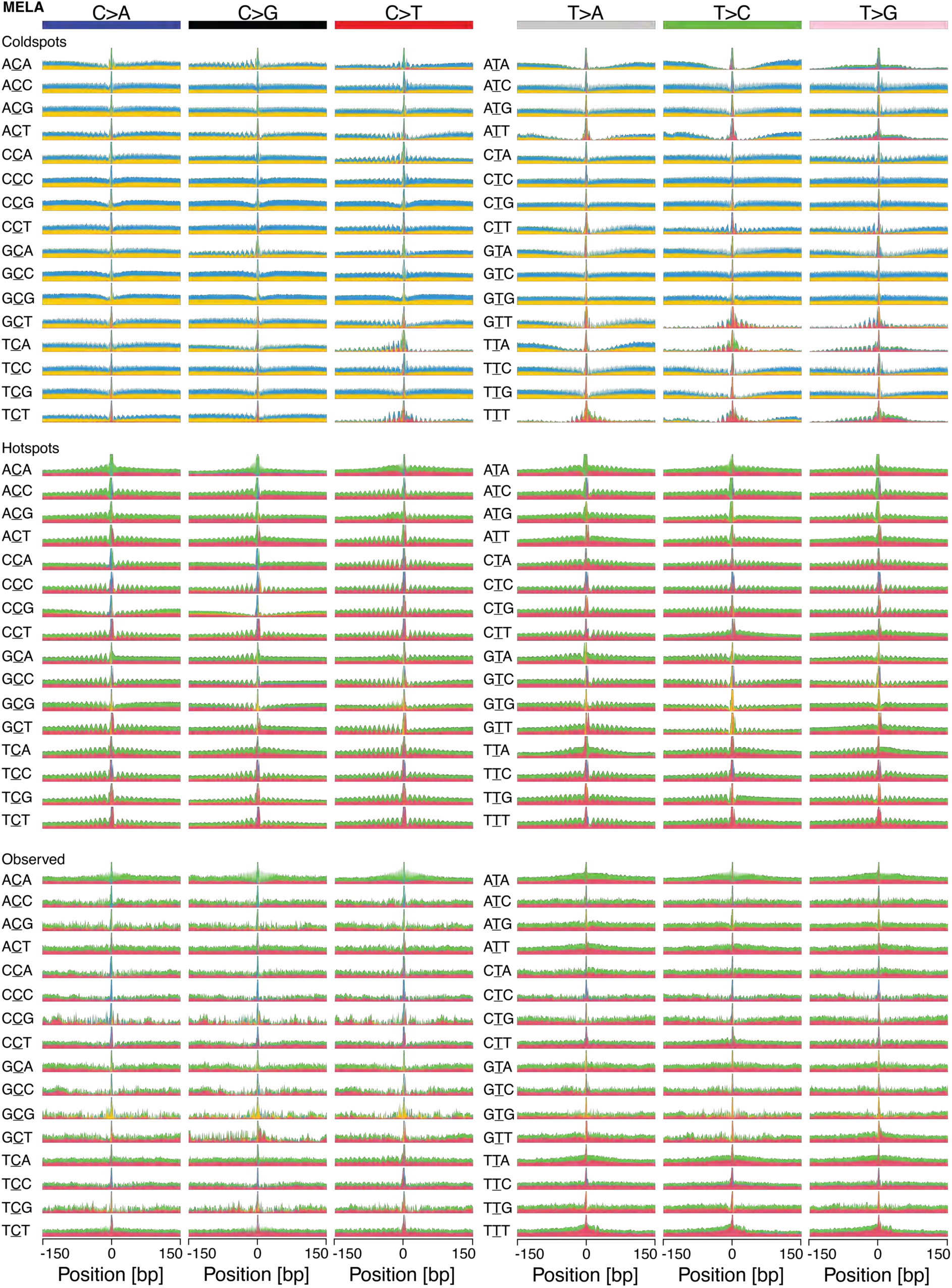
Mutation cold- and hotspots in MELA follow distinct intermediate range sequence motifs. Sequence motif patterns of 300bp for coldspots, hotspots and observed SBS, differentiated by their mutation type and trinucleotide context as before. Information content represents the differential of 2 bits and the information entropy of the sequences flanking the mutated trinucleotide, resolved by nucleotide as a motif visualization. The data correspond to the aggregated information content of FigureS10.

**Figure S15.**
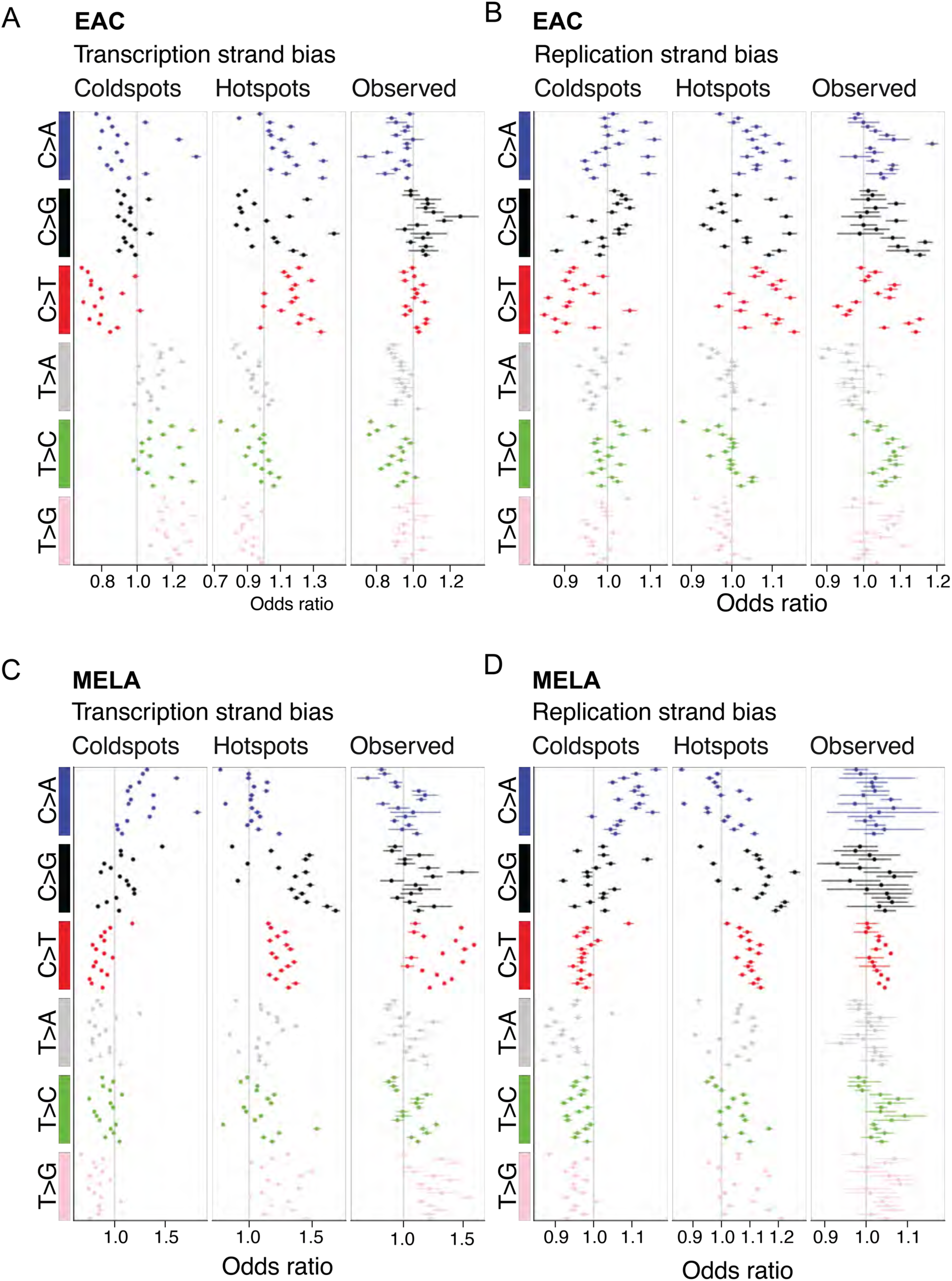
Mutation cold- and hotspots follow distinct strand bias. (A) Transcription strand bias for coldspots, hotspots and observed SBS in EAC, differentiated by their mutation type and trinucleotide context as before. Calculated are odds ratios with associated p-values based on the imbalance of the strands, corrected for the underlying bias of trinucleotide distribution. (B) DNA replication strand bias for coldspots, hotspots and observed SBS in EAC, differentiated by their mutation type and trinucleotide context as before. Calculated are odds ratios with associated p-values based on the imbalance of the strands, corrected for the underlying bias of trinucleotide distribution. (C) Transcription strand bias for coldspots, hotspots and observed SBS in MELA, differentiated by their mutation type and trinucleotide context as before. Calculated are odds ratios with associated p-values based on the imbalance of the strands, corrected for the underlying bias of trinucleotide distribution. (D) DNA replication strand bias for coldspots, hotspots and observed SBS in MELA, differentiated by their mutation type and trinucleotide context as before. Calculated are odds ratios with associated p-values based on the imbalance of the strands, corrected for the underlying bias of trinucleotide distribution

**Figure S16.**
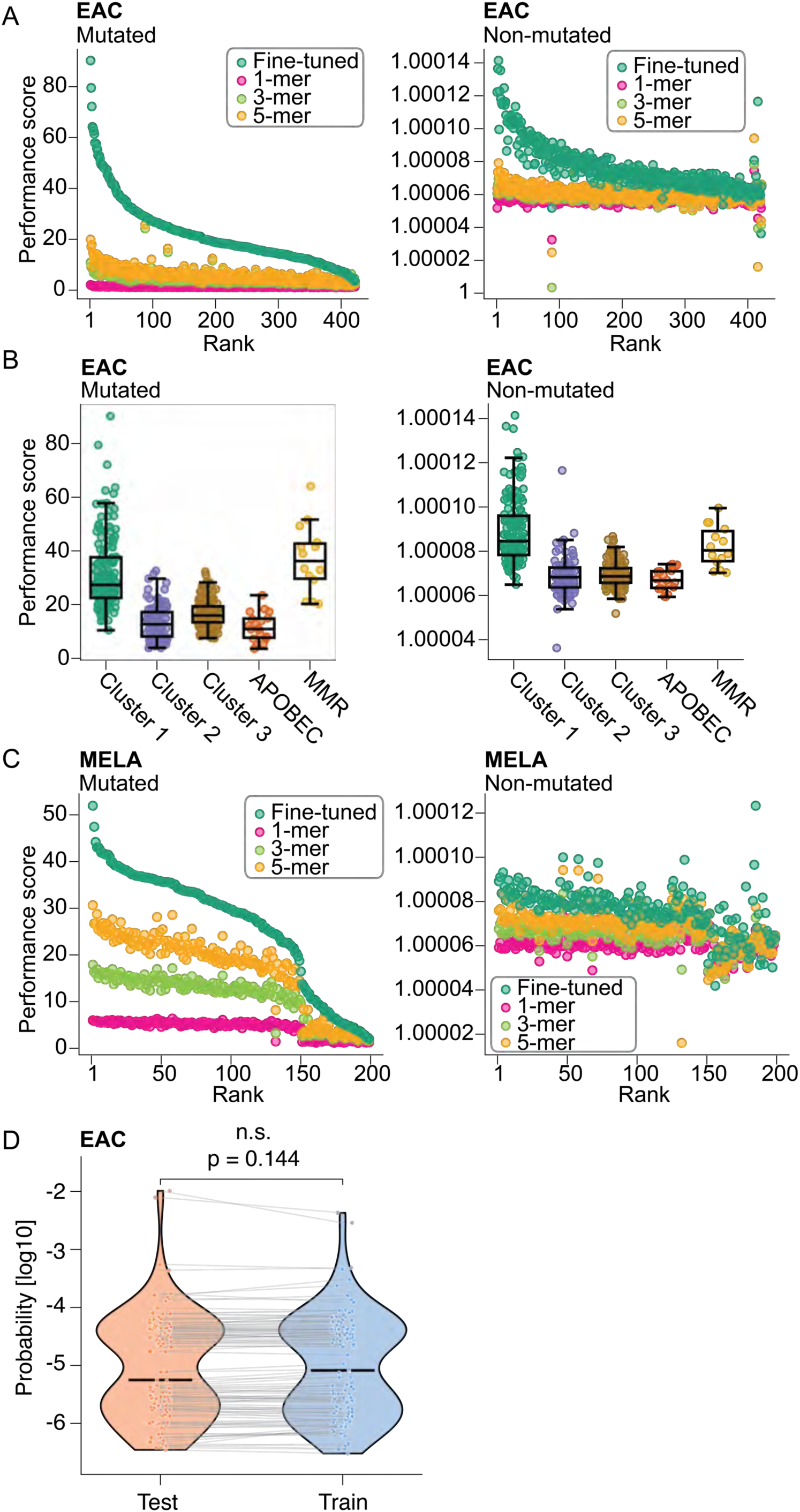
Fine-tuning of EAGLE per patient. (A) Performance assessment for individual samples after fine-tuning as the ratio of EAC EAGLE-predicted probability in the test set at mutated sites and non-mutated sites in comparison to proportional assignment of probabilities based on the observed mutations in the associated k-mer context. (B) Performance for individual samples differentiated by mutational subgroups. (C) Performance assessment for individual samples after fine-tuning as the ratio of MELA EAGLE-predicted probability in the test set at mutated sites and non-mutated sites in comparison to proportional assignment of probabilities based on the observed mutations in the associated k-mer context. (D) Predicted probability of 121 *TP53* mutations in EAC when the mutated position was in the test versus the training set of the patient’s own model (lines connect the same mutation; paired Wilcoxon signed-rank test).

**Figure S17.**
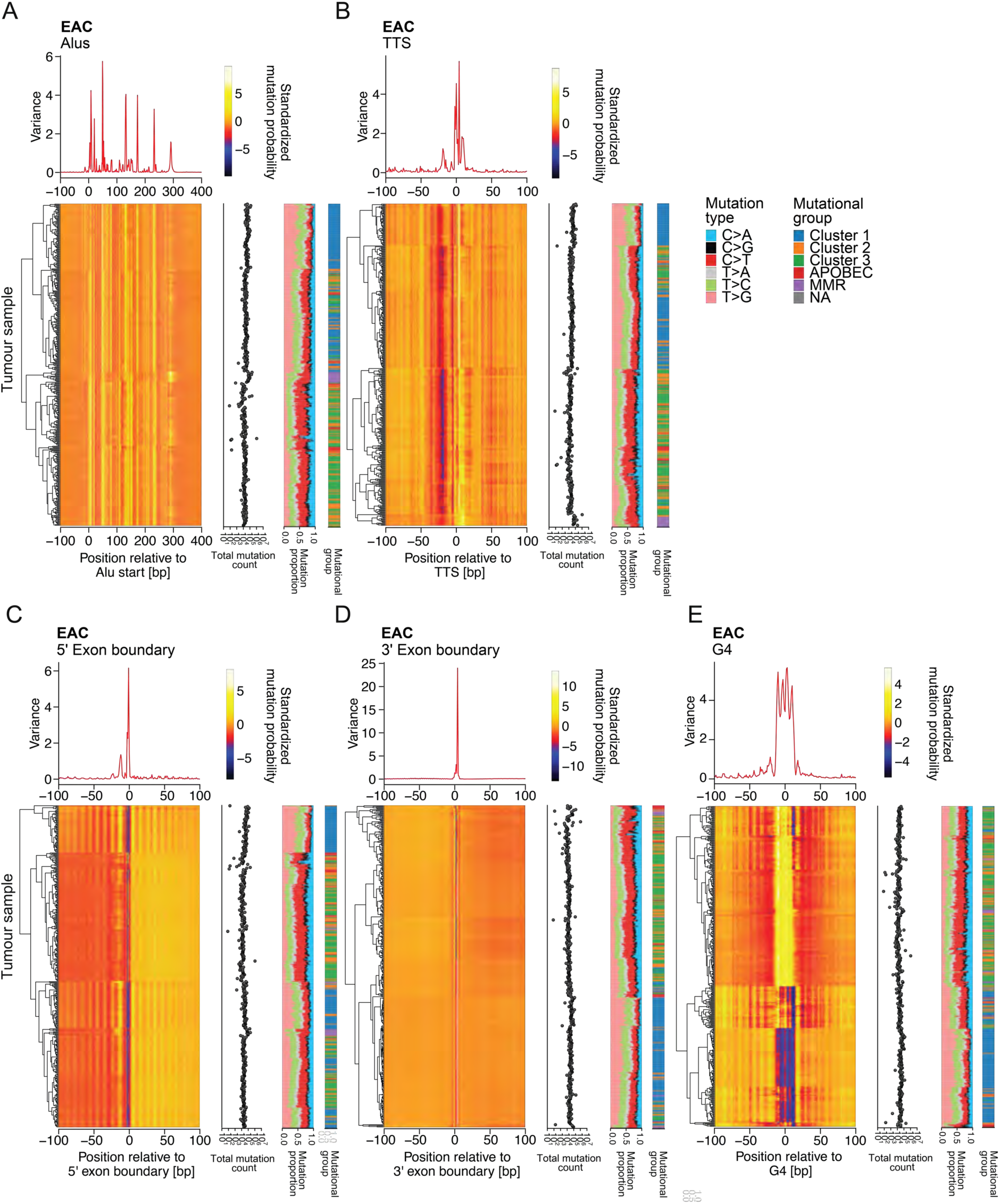
EAGLE predicted mutation probabilities per sample in EAC show individual differences of mutagenesis relative to genome-biological features. (A) Predictions of mutation probability relative to Alu retrotransposons (Alus) scaled by row. Samples are clustered by hierarchical clustering, set into relation with proportion of mutation types and mutational subgroups. Variance assessed per nucleotide on scaled probabilities. Analogously, data are presented for Transcriptional Termination Site (TTS) (B), 5’ exon boundary (C), 3’ exon boundary (D) and predicted G-quadruplex structures (G4s) (E).

**Figure S18.**
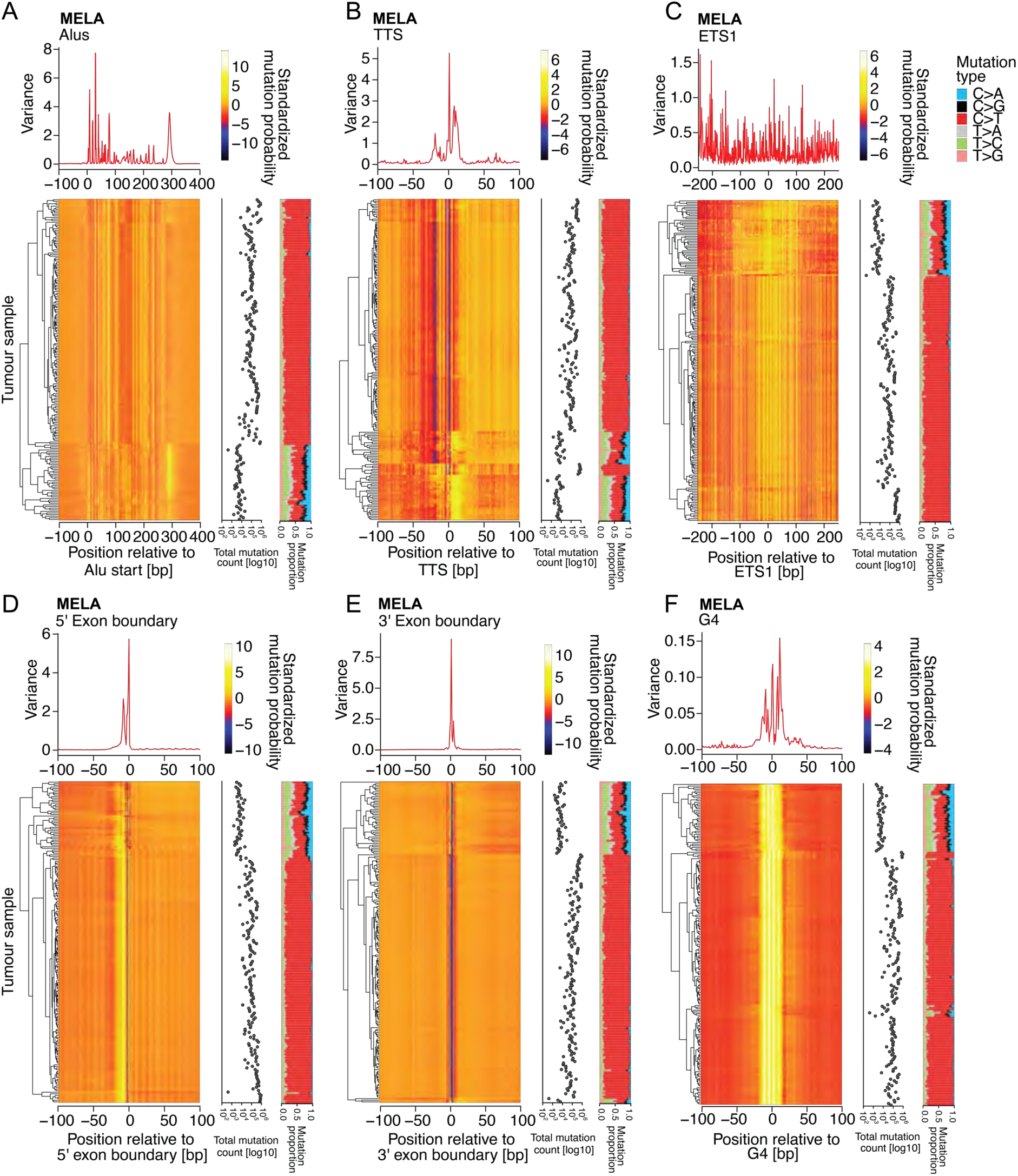
EAGLE predicted mutation probabilities per sample in MELA show individual differences of mutagenesis relative to genome-biological features. (A) Predictions of mutation probability relative to Alu retrotransposons (Alus) scaled by row. Samples are clustered by hierarchical clustering, set into relation with proportion of mutation types and mutational subgroups. Variance assessed per nucleotide on scaled probabilities. Analogously, data are presented for Transcriptional Termination Site (TTS) (B), ETS1 binding factor (C), 5’ exon boundary (D), 3’ exon boundary (E) and predicted G-quadruplex structures (G4s) (F).

**Figure S19.**
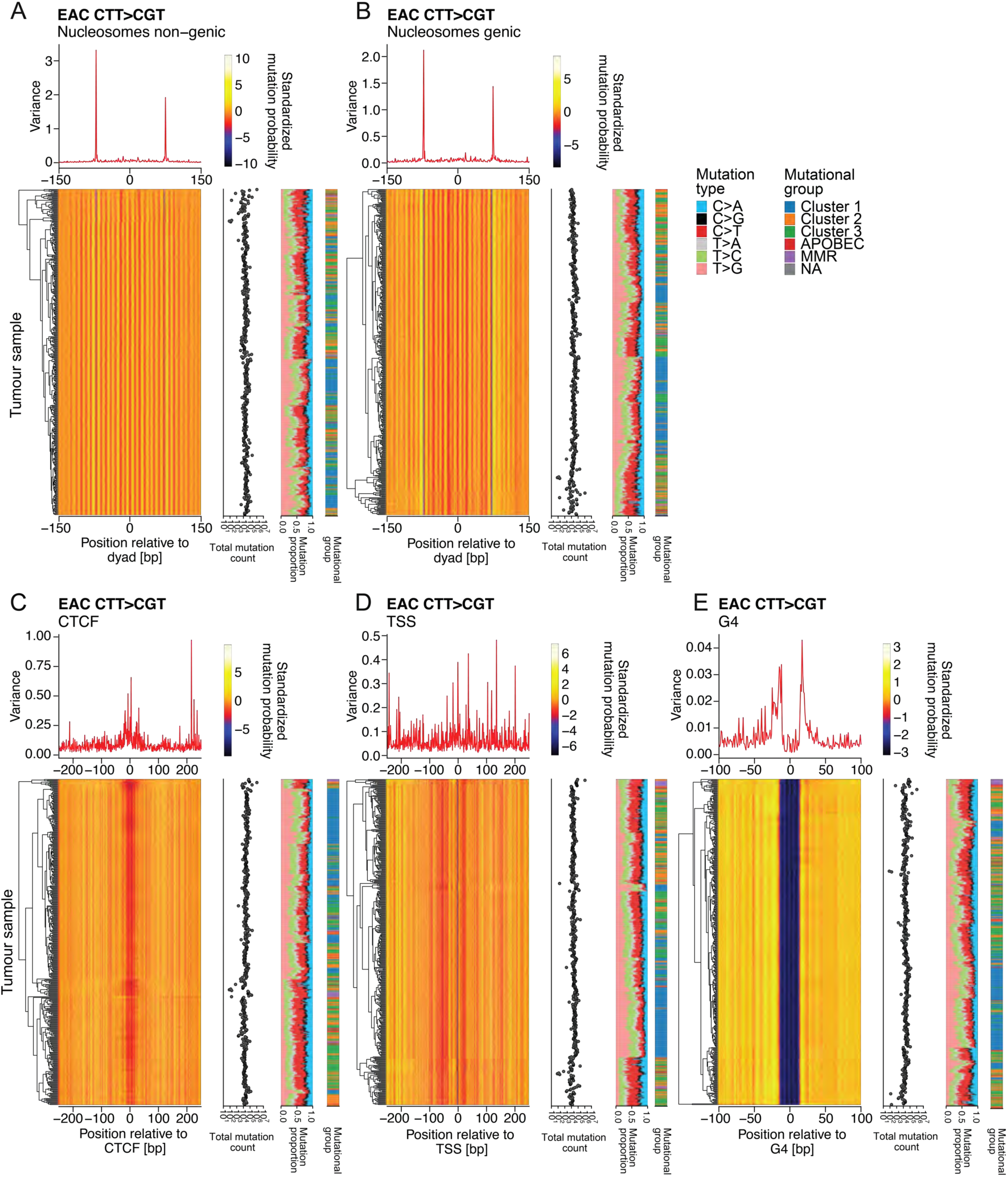
EAGLE predicted CTT>CGT/AAG>ACG mutation probabilities per sample in EAC. (A) Predictions of mutation probability of CTT>CGT/AAG>ACG relative to non-genic nucleosomes scaled by row. Samples are clustered by hierarchical clustering, set into relation with proportion of mutation types and mutational subgroups. Variance assessed per nucleotide on scaled probabilities. Analogously, data are presented for genic nucleosomes (B), CTCF = CCCTC-Binding Factor binding site (CTCF) (C), Transcriptional Start Site (TSS) (D), predicted G-quadruplex structures (G4s) (E).

**Figure S20.**
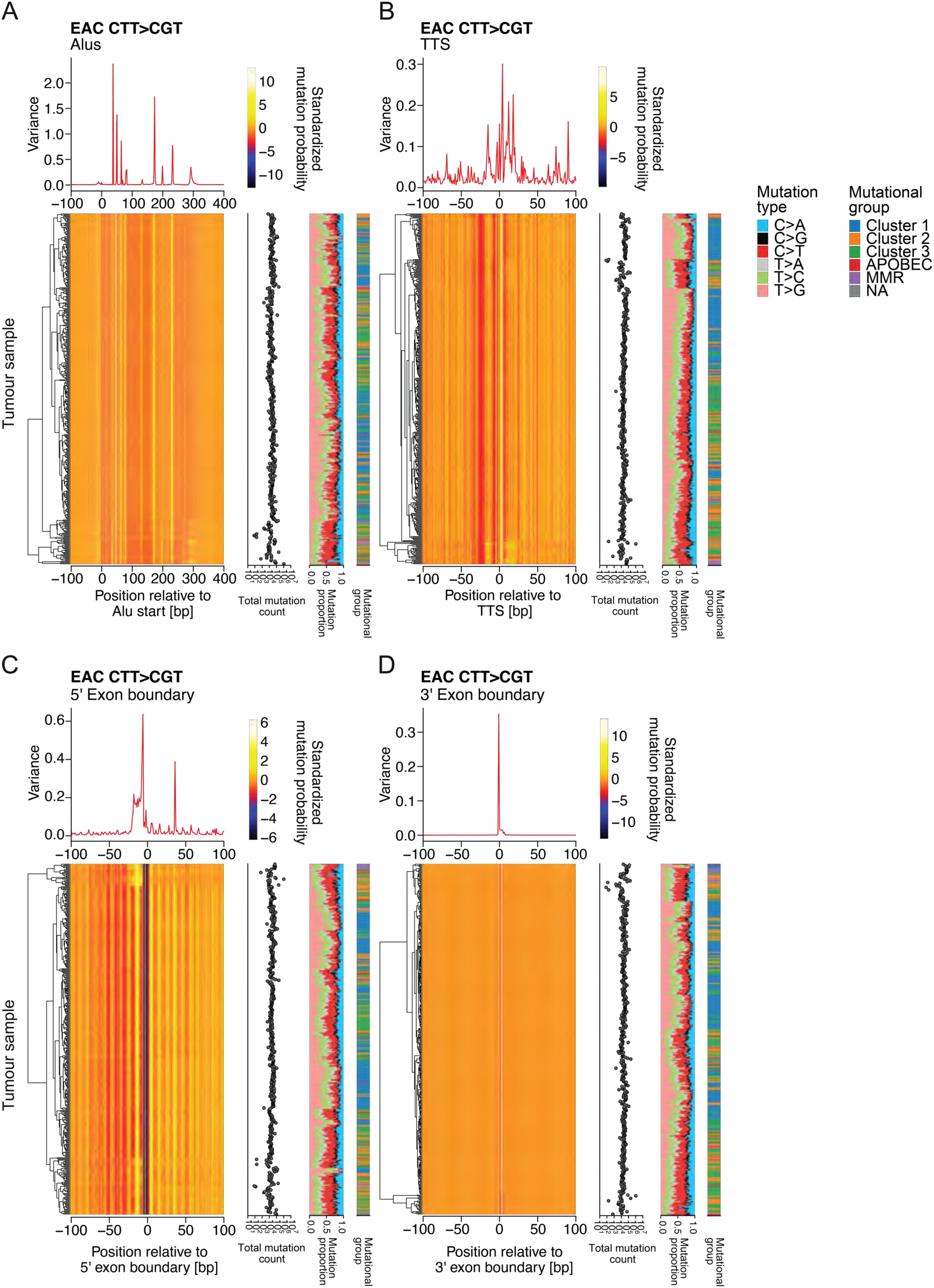
EAGLE predicted CTT>CGT/AAG>ACG mutation probabilities per sample in EAC. (A) Predictions of mutation probability of CTT>CGT/AAG>ACG relative to Alu retrotransposons (Alus) scaled by row. Samples are clustered by hierarchical clustering, set into relation with proportion of mutation types and mutational subgroups. Variance assessed per nucleotide on scaled probabilities. Analogously, data are presented for Transcriptional Termination Site (TTS) (B), 5’ exon boundary (C), and 3’ exon boundary (D).

**Figure S21.**
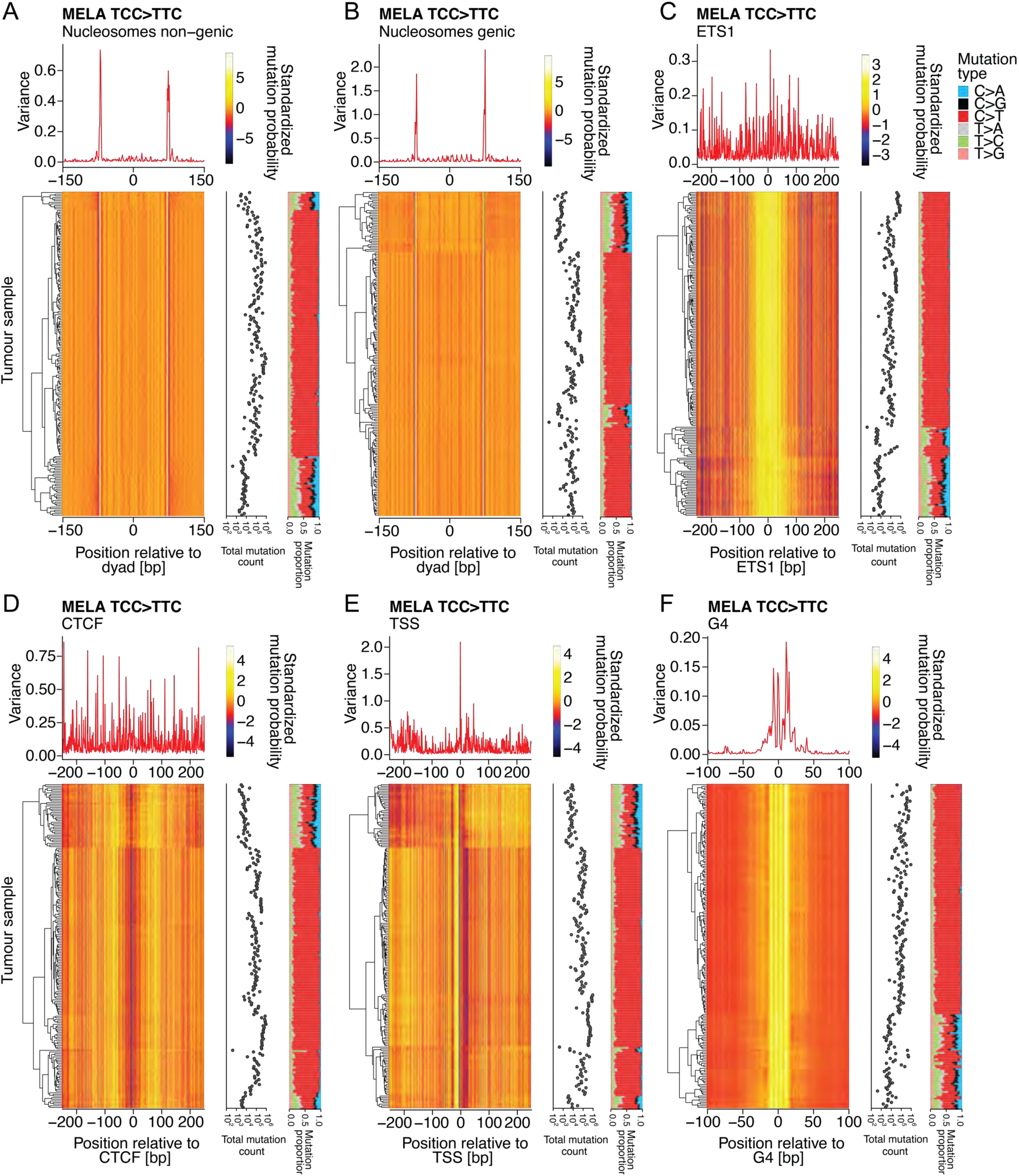
EAGLE predicted TCC>TTC/GGA>GAA mutation probabilities per sample in MELA. (A) Predictions of mutation probability of TCC>TTC/GGA>GAA relative to non-genic nucleosomes scaled by row. Samples are clustered by hierarchical clustering, set into relation with proportion of mutation types and mutational subgroups. Variance assessed per nucleotide on scaled probabilities. Analogously, data are presented for genic nucleosomes (B), ETS1 binding factor (C), CTCF = CCCTC-Binding Factor binding site (CTCF) (D), Transcriptional Start Site (TSS) (E), predicted G-quadruplex structures (G4s) (F).

**Figure S22.**
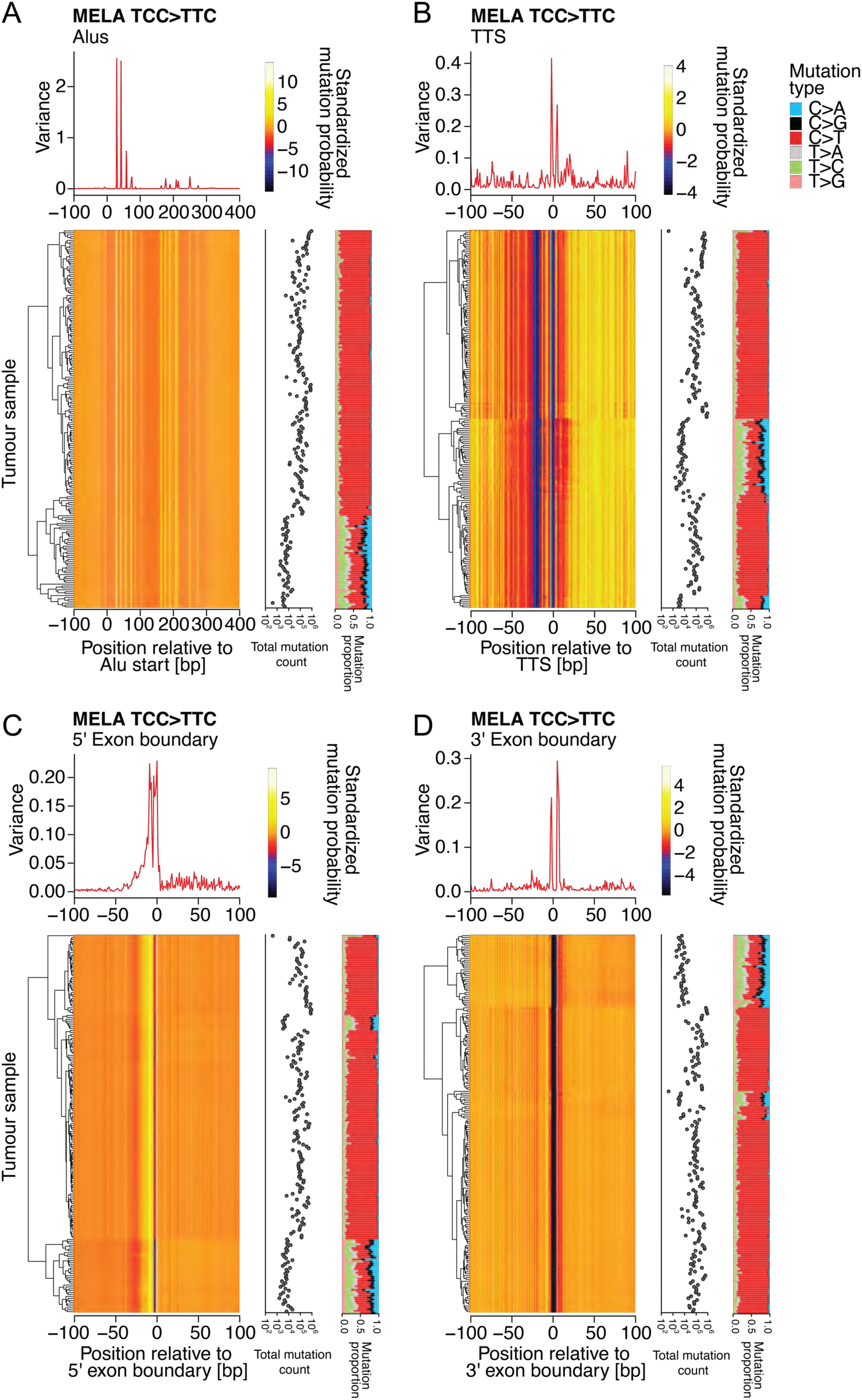
EAGLE predicted TCC>TTC/GGA>GAA mutation probabilities per sample in MELA. (A) Predictions of mutation probability of TCC>TTC/GGA>GAA relative to Alu retrotransposons (Alus) scaled by row. Samples are clustered by hierarchical clustering, set into relation with proportion of mutation types and mutational subgroups. Variance assessed per nucleotide on scaled probabilities. Analogously, data are presented for Transcriptional Termination Site (TTS) (B), 5’ exon boundary (C), and 3’ exon boundary (D).

